# Norovirus MLKL-like pore forming protein initiates programed cell death for viral egress

**DOI:** 10.1101/2023.03.17.533118

**Authors:** Guoxun Wang, Di Zhang, Robert Orchard, Dustin C. Hancks, Tiffany A. Reese

## Abstract

Non-enveloped viruses require cell lysis to release new virions from infected cells, suggesting that these viruses require mechanisms to induce cell death. Noroviruses are one such group of viruses, but a mechanism of norovirus-infection triggered cell death and lysis are unknown. Here we have identified a molecular mechanism of norovirus-induced cell death. We found that the norovirus-encoded NTPase contains a N-terminal four helix bundle domain homologous to the pore forming domain of the pseudokinase Mixed Lineage Kinase Domain-Like (MLKL). Norovirus NTPase acquired a mitochondrial localization signal, thereby inducing cell death by targeting mitochondria. NTPase full length (NTPase-FL) and N-terminal fragment (NTPase-NT) bound mitochondrial membrane lipid cardiolipin, permeabilized mitochondrial membrane and induced mitochondrial dysfunction. Both the N-terminal region and the mitochondrial localization motif of NTPase were essential for cell death, virus egress from cells and virus replication in mice. These findings suggest that noroviruses stole a MLKL-like pore forming domain and co-opted it to facilitate viral egress by inducing mitochondrial dysfunction.

## Main

Viruses mimic host factors to gain selective advantage over the host and subvert host defense strategies (*1*). Because programmed cell death is a critical host defense strategy against viral infection, many viruses have evolved strategies to evade or block programmed cell death (*2, 3*). On the other hand, some viruses require cellular lysis as the final step of virus replication, suggesting that viruses may have also evolved mechanisms to induce cell death.

Noroviruses are small, positive-sense, single-stranded RNA viruses of the family *Caliciviridae* (*4*–*6*). Norovirus genomes are organized into three open reading frames (ORFs), ranging in size from 7.3 to 7.5 kb. As non-envelope viruses, noroviruses are classically thought to egress from infected cells through cellular lysis (*7*–*9*), with some reports of exosomal release of virus *in vivo* (*10*). However, the requirement for particular cell death pathways is ill-defined and whether induction of cell death is required for newly assembled viruses to egress from the host cell is still unclear.

NINJ1 is required for end-stage lysis of plasma membrane in response to multiple types of cell death (*11*). To investigate whether NINJ1 is also involved in norovirus-induced cytotoxicity and viral egress, we infected wild-type and *Ninj1*^*-/-*^ BV2 cells, a mouse microglial cell line, with acute and persistent strains of murine norovirus (MNoV), CR6 and CW3, respectively. After infection, we observed that *Ninj1*^*-/-*^ cells swelled, developing bubble-like herniations, and retained this balloon morphology even 48 hours post infection (Fig. 1A). Consistent with previous data, the absence of NINJ1 did not prevent virus-triggered cell death, as measured by ATP levels (Fig. 1B) (*11*). However, NINJ1 deficiency inhibited lactate dehydrogenase (LDH) release in response to noroviruses infection, indicating that the plasma membrane remained intact (Fig. 1C). To determine if plasma membrane rupture was required for efficient virus release, we measured virus titer in the supernatant of infected cells 12 and 24 hours after virus infection. There was a marked decrease in virus detected in the supernatant of *Ninj1*^*-/-*^ cells compared with wildtype cells, even though wildtype and knockout cells produced equivalent amounts of intracellular virus (Fig. 1D). Together, these data suggest that NINJ1 mediates plasma membrane rupture during norovirus infection and regulates virus egress from infected cells.

**FIG. 1.**
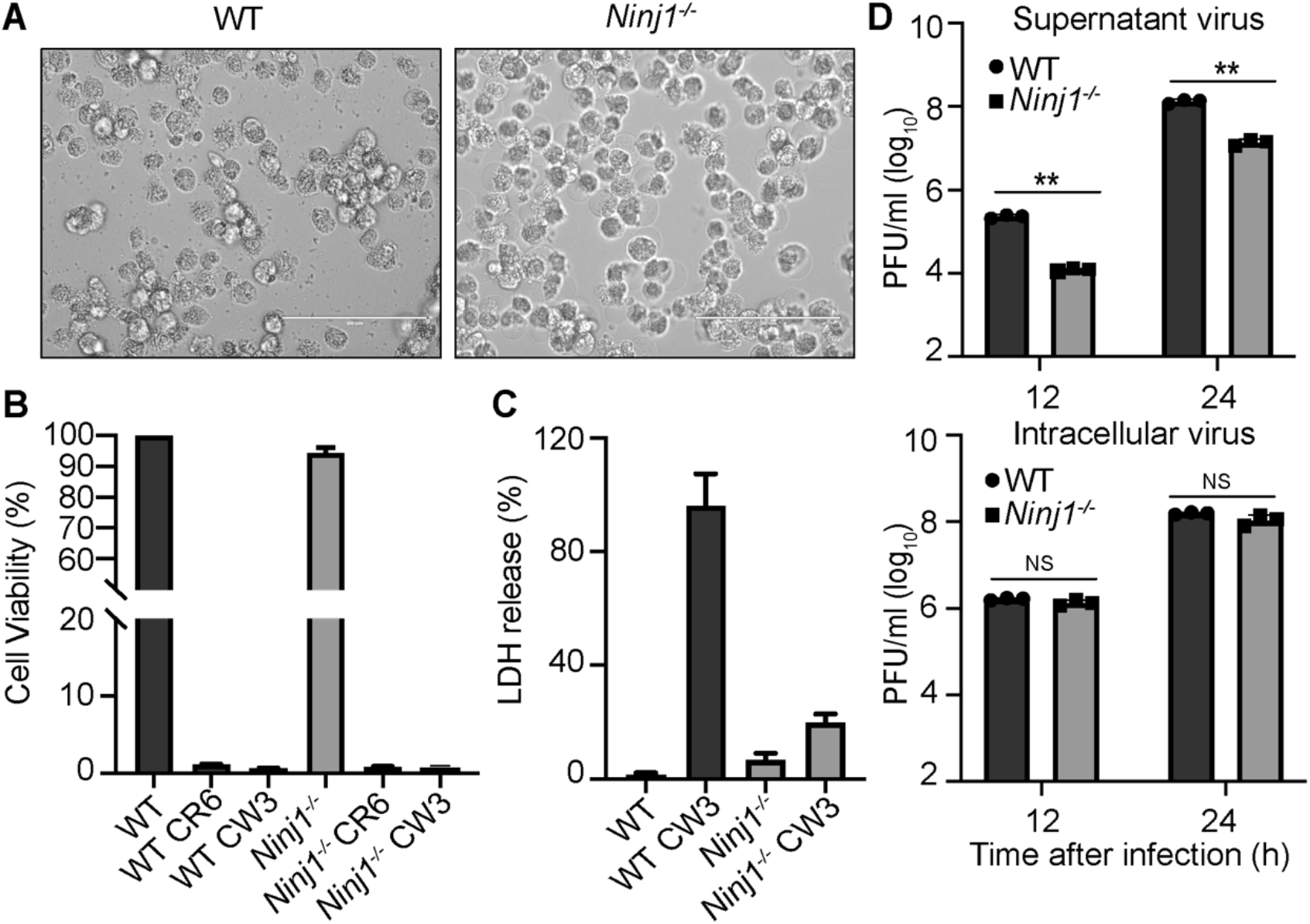
Norovirus egress requires NINJ1-mediated plasma membrane rupture. (**A**) Representative bright-field images of WT and *Ninj1*^*-/-*^ BV2 cells infected with MNoV^CR6^ at a MOI of 5 for 48 h. (**B**) Viability of WT and *Ninj1*^*-/-*^ BV2 cells infected with MNoV^CW3^and MNoV^CR6^ at a MOI of 5 for 24 h. (**C**) LDH released from WT and *Ninj1*^*-/-*^ BV2 infected with MNoV^CR6^ at a MOI of 5 for 12 h. (**D**) WT and *Ninj1*^*-/-*^ BV2 cells infected with MNoV^CR6^ at a MOI = 1 and supernatant or cell associated (intracellular) virus was measured by plaque assay at indicated time. Data represented in (**B, C** and **D)** are expressed as mean values ± s.d. from three technical replicates. Data are representative of three independent experiments. Statistical analysis was conducted using two-way ANOVA followed by Tukey’s multiple comparison test, NS, not significant; *P<0.05; **P<0.01; ***P<0.001.

Because NINJ1 is downstream of multiple programmed cell death pathways, we next asked which upstream cell death pathways are required for norovirus-induced cell lysis. To characterize the cytotoxicity of noroviruses infection, we infected either bone marrow-derived macrophages (BMDMs) or BV2 cells with MNoV strains CW3 and CR6 (*12*). We observed a dramatic decrease in the viability of both cell types and detected LDH release into the culture medium at 24 hours after infection (Fig. S1A, B and S2). Infected cells swelled and stained positive for annexin V and propidium iodide at a late timepoint post infection (Fig. S1C). To determine whether necroptosis, pyroptosis and/or apoptosis were required for cell death, we added the necroptosis inhibitors, necrostatin-1 or MLKLi, and the pan-caspase inhibitor z-VAD during MNoV infection to block programmed cell death (*13, 14*). None of these treatments, either individually or in combination, were able to block noroviruses-induced cell death. These data suggest that key effectors of necroptosis, pyroptosis and apoptosis are not required for MNoV-induced cell death (Fig. S3A-D and S4).

Taking a genetic approach, we assessed cell death in a variety of knockout cells. To evaluate pyroptosis, we infected macrophages isolated from *Caspase1/11*-deficient mice and BV2 cells deficient in *Gasdermin D* (*Gsdmd)* with MNoV. For infected wildtype and knockout cells, similar decreases in cell viability were observed, even in the presence of necroptosis inhibitors, indicating that pyroptosis-induced cell death is dispensable (Fig. S5A-D). To assess necroptosis, BMDMs were infected with CR6 or CW3 strains of MNoV and phosphorylation of MLKL, which is a hallmark of necroptotic signaling, was measured (*15*). Neither MNoV strain induced phosphorylation of MLKL (Fig. S6A). Furthermore, macrophages from *Ripk3*-deficient mice or BV2 cells deficient in *Mlkl*, had a comparable loss ian cell viability after MNoV infection (Fig. S6B-D). Treating *Mlkl*-deficient cells with zVAD could not rescue cell death, suggesting that necroptosis is also dispensable (Fig. S6B-D). Finally, to evaluate apoptosis, we generated BV2 *Bax/Bak* double knockout, *Casp9/Gsdmd* double knockout, and *Casp3/Gsdmd* double knockout cells and measured viability after MNoV infection. All knockout cells exhibited cell death equivalent to wildtype cells after MNoV infection (Fig. S7A-D). Unexpectedly, these data indicate that MNoV-induced cell death does not require host executioners of pyroptosis, necroptosis, or apoptosis.

Notably, we observed cleavage of CASPASE-3 and GSDMD in infected cells (Fig. S8A) (*7*–*9*). Although neither *Caspase-3* nor *Gsdmd* were required for norovirus-induced cell death, these data indicate that MNoV infection initiates programmed cell death, which may trigger secondary necrosis. We next investigated whether a norovirus-encoded protein could directly induce cell death.

To determine whether viral proteins directly trigger programmed cell death, we ectopically expressed individual MNoV genes in human embryonic kidney (HEK)-293T cells. Expression of NTPase (NS3), but not other viral genes, induced marked cytotoxicity based on ATP levels and LDH release, which was not blocked by the pan-caspase inhibitor, zVAD (Fig. 2A, Fig. S9A-E and Fig. S10C). Norovirus NS3 is a highly conserved nonstructural protein that displays NTPase activity, which is essential for viral genome synthesis. Norovirus NS3 may also promote apoptosis in transfected cells (*16, 17*). NS3 comprises three domains: The N-terminal domain (NTD, 1-158), the core domain (158–289) and the C-terminal domain (CTD, 290–364) (*18, 19*). The evolutionarily conserved core domain and the C-terminal domain, hereafter called NTPase-CT, contain the evolutionarily conserved viral helicase and peptidase, which is required for viral genome replication (*18, 20*). However, like many other small RNA viruses, norovirus proteins often perform multiple functions in the replicative cycle (*6, 21, 22*). The N-terminal domain, NTPase-NT has no known function in the viral life cycle.

**FIG. 2.**
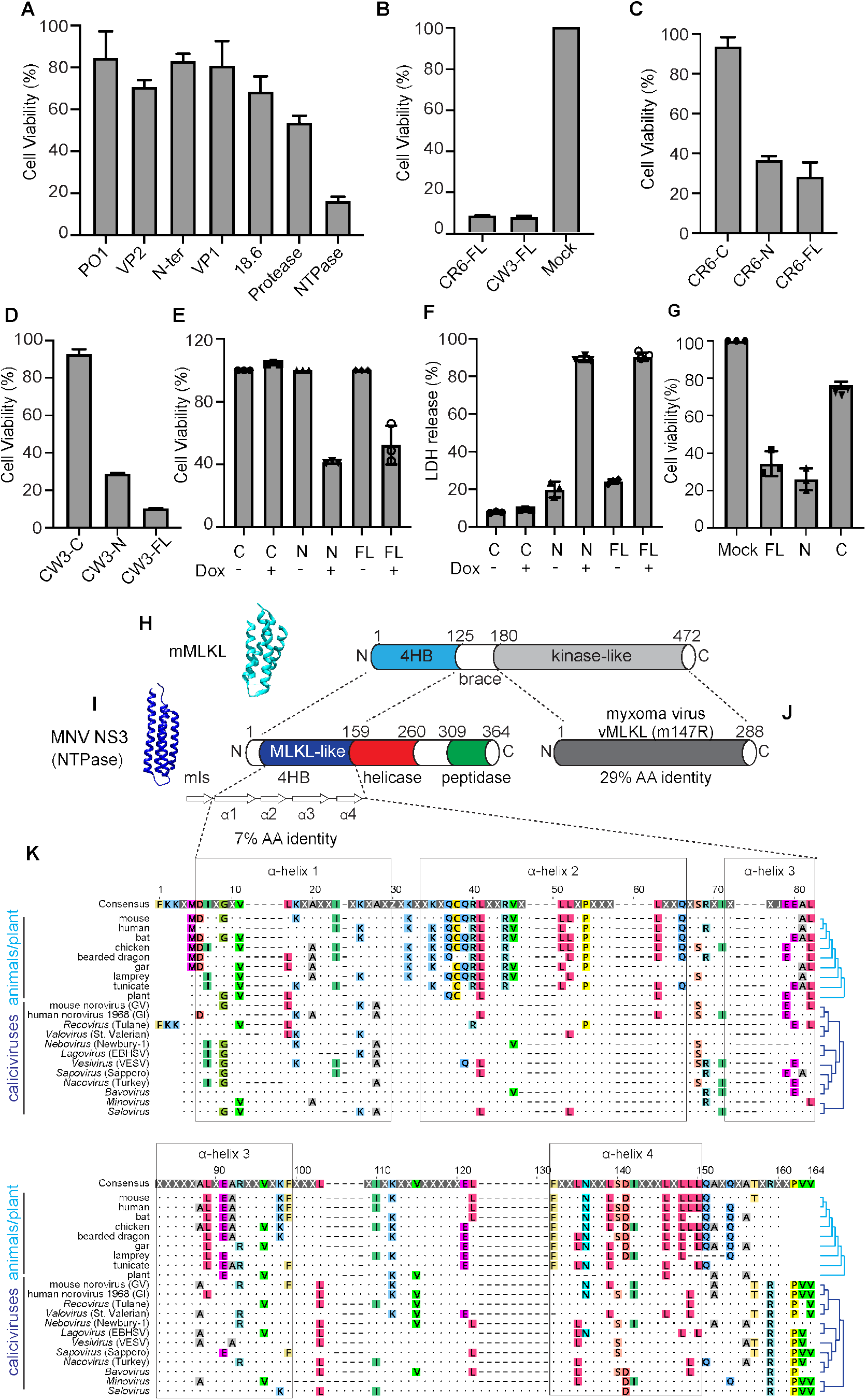
N-terminal domain of norovirus NTPase directly triggers programmed cell death and has a 4-helix bundle domain with homology to MLKL. (**A**) The genes of MNoV strains CR6 were transfected into HEK293T cells. Cell death was determined by ATP-based cell viability assay 18 h after transfection. (**B**) Cell death was determined by ATP-based cell viability assay 18 h after transfection of NTPase of MNoV strains CR6 and CW3. (**C**) Full-length (F), N- or C-terminal of NTPase from MNoV strain CR6 was transfected into HEK293T cells. Cell death was determined by ATP-based cell viability assay 18 h after transfection. (**D**) Full-length (F), N- or C-terminal of NTPase from MNoV strain CW3 was transfected into HEK293T cells. Cell death was determined by ATP-based cell viability assay 18 h after transfection. (**E-F**) Full-length, N- or C-terminal of MNoV strains CR6 NTPase were stably expressed in BV2 cells under a tetracycline-inducible promoter. Transgene expression was induced with doxycycline treatment. ATP-based cell viability (**E**) and LDH release (**F**) analysis after 12 h doxycycline induction. (**G**) Effects of HNoV MD145 NTPase on cell death activity. Full-length, N- or C-terminal of NTPase were transfected into 293T cells. Cell death was determined by ATP-based cell viability assay 18 h after transfection. Data represented in (**A**-**G**) are expressed as mean values ± s.d. from three technical replicates. Data are representative of three independent experiments. (**H**) Model of 4HB for mouse MLKL (PDB: 4BTF) and protein schematic (*38*). (**I**) Model of N-terminal 4HB for MNV NS3 and protein schematic. Model generated using Robetta (https://robetta.bakerlab.org/) and 4HB from mouse MLKL (PDB: 4BTF); Visualized using Chimera (https://www.cgl.ucsf.edu/chimera/). Secondary structure predicted using JPred4 (http://www.compbio.dundee.ac.uk/jpred/). α-helices are displayed as white, right facing arrows. Helicase and peptidase domain annotations based on NCBI conserved domains (https://www.ncbi.nlm.nih.gov/Structure/cdd/wrpsb.cgi). mls: putative mitochondrial localization signal. (**J**) Poxviruses encode a MLKL-mimic derived from the kinase-like domain (*34, 35*). (**K**) Amino acid multiple sequence alignment for 4HB domains generated using MUSCLE with representative sequences: animal (8 species), plant (AtMLKL3)(*39*), and calicivirus genera (12 species; ICTV and NCBI). Helix boundaries are derived from the mouse MLKL predicted structure. Top line displays consensus. Conserved amino acid residues are individually colored. X and dots indicate hypervariable sites. GV - genogroup V; GI – genogroup I; EBHSV – European brown hare syndrome virus; VESV - vesicular exanthema of swine virus. Expanded alignments and accession numbers present in supplemental tables 1-3.

To gain insights into this NS3-associated cell death activity, we performed protein-modeling and detailed sequence analysis. Previous protein modeling suggested that the N-terminus of MNV NS3 may encode a four-helix bundle domain (4HB) (*18, 20*). Because the executioner of necroptosis, MLKL, also encodes a 4HB domain, we hypothesized that the N-terminus of MNV NS3 may encode for a MLKL-like protein. To examine this, we predicted the secondary structure for residues 1-158 of MNV NS3, because NCBI Conserved Domain Database predicts the RNA helicase starts at residue 159. JPred4 predicted five α-helices within residues 1-158 (Fig. 2G). To determine whether the MNV NS3 N-terminal domain resembles the 4HB domain of mouse MLKL, we performed protein modeling for NS3 residues 25-158 with the published structure of the mouse protein (PDB: 4BTF) (Fig. 2H). NS3 residues 1-24 were excluded given the predicted mitochondrial targeting sequence residing in the first putative α-helix (Fig. S11). Consistently, modeling of the N-terminus of MNV NS3 shows that this domain may form a 4HB domain. To further understand the relationship between the 4HB domain of cellular MLKL proteins and NS3, we analyzed thirty-four divergent cellular MLKL sequences from animals and plants with thirty-one calicivirus sequences across known viral genera, including representatives from established norovirus genogroups (supplemental tables 1-3). This analysis (Fig. 2I and Fig. S12) revealed residual sequence identity, which overlaps with the α-helices, between these cellular and viral 4HB domains even for the most divergent *caliciviruses*, such as *saloviruses*. These data suggest that the N-terminus of *caliciviruses* NS3 encodes a MLKL-like protein.

Since the four-helix bundle domain and Helo-like domain have cell death-execution function, we hypothesized that the N-terminal domain of NTPase induces cell death (*23, 24*). As predicted, expression of NTPase-NT from CR6 and CW3 induced cell death, but expression of NTPase-CT did not induce appreciable cell death (Fig. 2B). We next tested whether the NTPase of a human norovirus strain (HNoV), MD145, also promoted cell death. Similar to MNoV NTPase, HNoV NTPase induced cell death in 293T cells overexpressing full length or the N-terminal fragment of MD145 HNoV NTPase, but not the C-terminal fragment (Fig. 2C, 2G and Fig. S13A-C) (*25*). These results indicate that the N-terminal domain of MNoV and a HNoV NTPase directly triggers programmed cell death.

To verify the role of NTPase-NT in programed cell death, we generated BV2 cells that stably express doxycycline (Dox)-inducible full length NTPase (NTPase-FL), NTPase-NT or NTPase-CT. Induction of NTPase-FL and NTPase-NT, but not NTPase-CT, resulted in cell lysis as measured by reduced ATP levels, release of LDH and uptake of SYTOX green (Fig. 2D and Fig. S10B, D and E). We further noted that the dying cells displayed morphological changes similar to norovirus infected cells, including cell rounding, swelling and plasma membrane rupture (Fig. S13A). These observations are consistent with the hypothesis that NTPase-NT directly drives cell death during norovirus infection.

Given that MLKL localizes to the plasma membrane (*23, 26*), we asked where NTPase-NT localized. DeepLoc analysis, which predicts subcellular localization for multiple compartments, indicated that MNV NS3 may have a mitochondrial localization signal at its immediate N-terminus (Fig. S14A, B) (*27*). To determine subcellular localization, we fractionated the post-nuclear supernatant of Dox-induced cell lines expressing NTPase-FL, NTPase-NT and NTPase-CT. While NTPase-CT was only detected in the cytosolic fraction, NTPase-FL and NTPase-NT were enriched in the mitochondrial fraction (Fig. 3A). To further visualize the subcellular distribution of NTPase, we examined the localization of the NTPase by immunofluorescence microscopy. While NTPase-CT was diffuse throughout the cytosol, NTPase-FL and NTPase-NT colocalized mitochondria marker, Cox IV (Fig. 3B). In contrast to MLKL, neither NTPase-FL nor NTPase-NT colocalized with the plasma membrane marker, Cell Mask Orange, indicating NTPase-FL and NTPase-NT primarily target mitochondria for executing cell death (Fig. S15) (*23*). Accordingly, cells expressing mutant forms of NTPase-FL and NTPase-NT lacking mitochondria localization motif displayed no cytotoxicity (Fig. 3C).

**FIG. 3.**
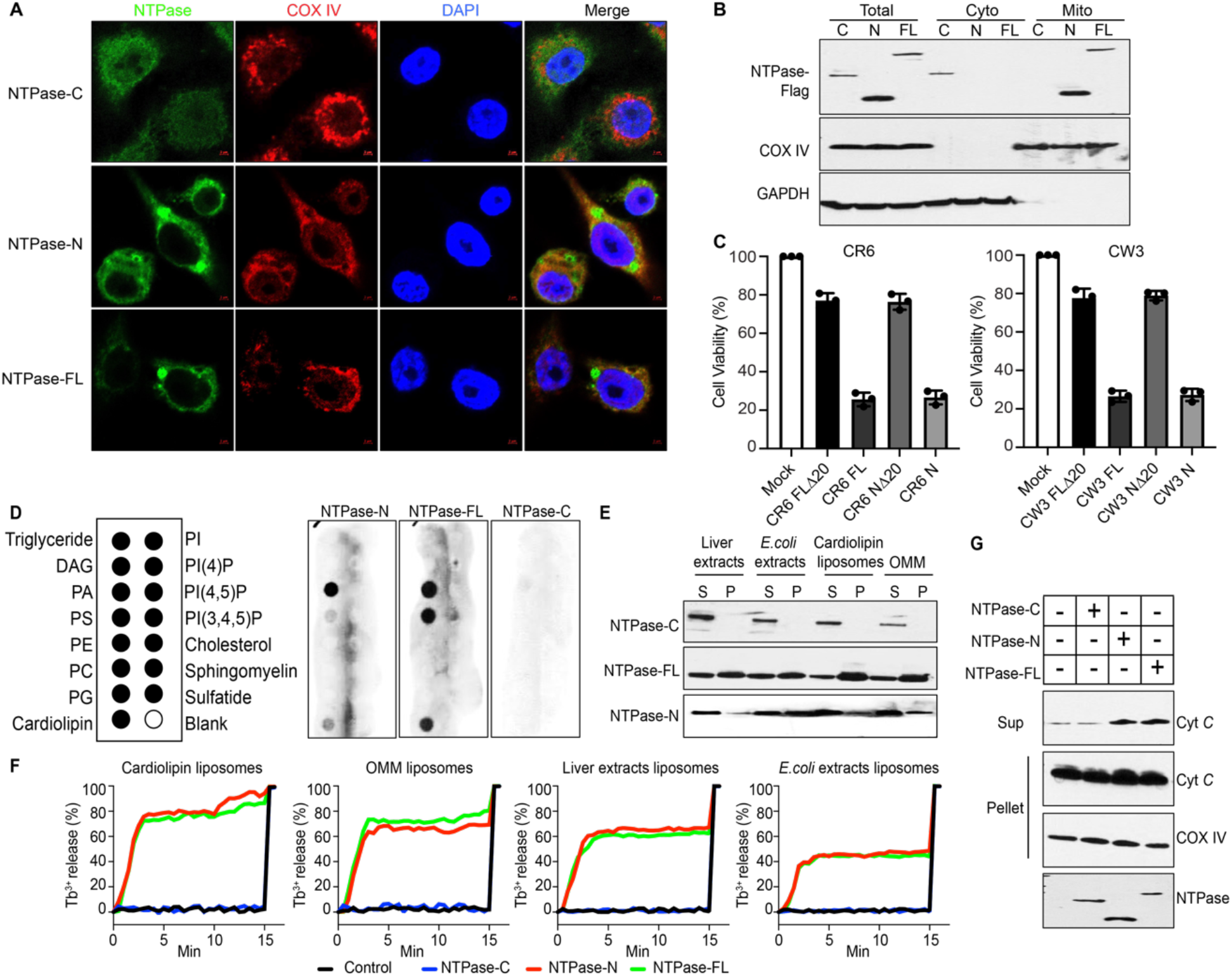
Norovirus NTPase permeabilizes the mitochondrial membrane and induces mitochondrial disfunction. (**A**) Representative confocal imaging of 293T cells after transfection with full-length, N- or C-terminal NTPase. The green channel shows the expression of NTPase. The red channel shows mitochondrial staining with cytochrome c oxidase IV (Cox IV). The merged channels are shown on the right. Scale bar, 10 µm. (**B**) Full-length, N- or C-terminal NTPase were stably expressed in BV2 cells under a tetracycline-inducible promoter. Cells were separated into cytosol and mitochondrial fractions after doxycycline induction and analyzed by immunoblot with indicated antibodies. (**C**) MNoV strains CR6 and CW3 NTPase full-length, N-terminal, full-length 20 (FL20), or N-terminal 20 (N20) mutants lacking the predicted mitochondrial localization signal were transfected into HEK293T cells. Cell viability was determined by ATP assay 18 h after transfection. (**D**) Purified full-length, N- or C-terminal NTPase proteins were incubated with lipid strips. Anti-MBP antibodies were used to detect the NTPase proteins on lipid strips. (**E**) Purified full-length, N- or C-terminal NTPase proteins were incubated with indicated liposomes. After ultracentrifugation, the liposome-free supernatant (S) and the liposome pellet (P) were analyzed by immunoblot. (**F**) Purified full-length, N- or C-terminal NTPase proteins were incubated with indicated liposomes. Liposome leakage was monitored by measuring DPA chelating-induced fluorescence of released Tb^3+^ relative to that of Triton X-100 treatment. Cardiolipin liposome: 80% PC + 20% cardiolipin; OMM liposome: 46% PC +25% PE + 11% PI + 10% PS + 7% cardiolipin. (**G**) Cyt *c* released into the reaction supernatant (Sup) from purified mitochondria (Mito) incubated with purified NTPase proteins. Data shown are representative of at least three (**B**-**G**) or two (**A**) independent experiments.

Given that NTPase-FL and NTPase-NT localize to mitochondria, we next examined whether the NTPase altered mitochondria membrane potential. To determine if MNoV NTPase depolarized mitochondria, cells expressing inducible NTPase constructs were stained with the cell permeate dye, TMRM, which is sequestered by active mitochondria. Upon loss of mitochondrial membrane potential, TMRM fluorescence decreases. Induction of NTPase-FL or NTPase-NT expression, but not NTPase-CT, led to depolarization of mitochondria as evidenced by reduced TMRM staining (Fig. S16, D and E). To detect mitochondrial reactive oxygen species, cells were incubated with MitoSOX after dox-induction of NTPase transgene expression. NTPase-FL and NTPase-NT, but not NTPase-CT expression, increased mitochondrial ROS abundance (Fig. S16C). MNoV infection also induced mitochondria ROS and impaired mitochondria membrane potential (Fig. S16A, B). Therefore, we conclude that the N-terminus of NTPase-NT promotes mitochondrial dysfunction and depolarization. Consistent with this notion, induction of NTPase-FL and NTPase-NT expression also triggered cleavage of CASPASE-3 (Fig. S8B), which suggests that NTPase induces mitochondrial depolarization.

We hypothesized that NTPase-NT disrupts mitochondrial outer membranes to initiate cell death. To test this hypothesis, we incubated recombinant NTPase-FL, NTPase-NT and NTPase-CT proteins with membrane lipid strips to determine if they interact with lipids found in different membranes of the cell. NTPase and NTPase-NT bound to phosphatidylserine (PS), phosphatidic acid (PA) and cardiolipin (CL), which are mitochondrial and bacterial lipids, but not to other phosphoinositides found in the plasma membrane. NTPase-CT did not bind to any lipids (Fig. 3D). We tested whether NTPase-NT could interact with liposomes containing different compositions of lipids. NTPase-NT and NTPase-FL, but not NTPase-CT, efficiently precipitated with liposomes containing cardiolipin and the carrier lipid PC, as well as liposomes from bovine liver extracts and *E. coli* polar extracts (Fig. 3E). We also observed interaction of NTPase-NT and NTPase-FL, but not NTPase-CT, with liposomes that mimic the outer membrane of mitochondria (OMM) (Fig. 3E). We determined whether NTPase-FL and NTPase-NT induced liposome disruption. Both NTPase-NT and NTPase-FL caused nearly 70% leakage of cardiolipin liposomes and OMM liposomes (Fig. 3F). Similar results were obtained when the NTPase-FL and NT proteins were incubated with liposomes generated from bovine liver extracts and *E. coli* polar extracts. They had no effect on PC-reconstituted liposomes (Fig. S17). On the other hand, NTPase-CT did not disrupt any liposomes (Fig. 3F). Consistent with the high binding affinity to cardiolipin and disruption of the liposome reconstituted from *E. coli* polar extracts, NTPase-FL and NTPase-NT are also toxic to *Escherichia coli* (Fig. S18A, B).

Because other cell death executioners like MLKL are hypothesized to form oligomers that insert into membranes, we next tested whether NTPase formed higher order oligomers. We con-transfected HEK293T cells with Flag and GFP-tagged NTPase-FL and NTPase-NT. Immunoprecipitation with anti-FLAG antibody readily pulled down the GFP-tagged NTPase-FL and NTPase-NT, indicating that NTPase-FL and NTPase-NT can self-associated and form higher order oligomers (Fig. S19).

To provide direct evidence that NTPase permeabilizes the mitochondrial membrane, we incubated mitochondria with purified NTPase proteins. Both NTPase-FL and NTPase-NT strongly induced cytochrome *c* release from the mitochondria, whereas NTPase-CT did not (Fig. 3G), These data suggest that the N-terminal domain of MNoV NTPase disrupts mitochondria by forming pores in the outer mitochondrial membrane by a process that potentially involves assembly of higher order oligomers.

The finding that NTPase-NT is sufficient to disrupt the mitochondrial membrane and induce programmed cell death, raised the possibility that NTPase-NT may be required for viral egress from infected cells. To determine whether NTPase-induced cell death was required for viral egress, we used the reverse genetics system for MNoV to generate mutant viruses lacking the N-terminal region of NTPase (MNoV^CW3^ΔN and MNoV^CR6^ΔN) or lacking mitochondrial localization motif in the NTPase (MNoV^CW3^ΔN20 and MNoV^CR6^ΔN20) in the two MNoV strains, CR6 and CW3 (*28*). In contrast with wildtype MNoV, MNoV^CW3^ΔN and MNoV^CR6^ΔN were unable to lyse BV2 cells, despite expressing viral non-structural proteins and forming replication complexes (Fig. 4A-C and Fig. S20A, B). By measuring viral genome copies that were cell associated or in the supernatant, we found that N-terminal truncation mutant viruses lost the capacity to egress the cell even though the mutant viruses replicated to comparable levels (Fig. 4D, E). Similarly, the mutant viruses lacking the mitochondrial localization motif (MNoV^CW3^ΔN20 and MNoV^CR6^ΔN20) did not induce cytotoxicity and were defective in viral release into the cell supernatant (Fig. S21A). The impaired viral egress was consistent with our difficulty obtaining high titer mutant viral stocks, necessitating the use of inducible complimenting cell lines to produce infectious virus. These results indicate that NTPase-NT is required for viral egress.

**FIG. 4.**
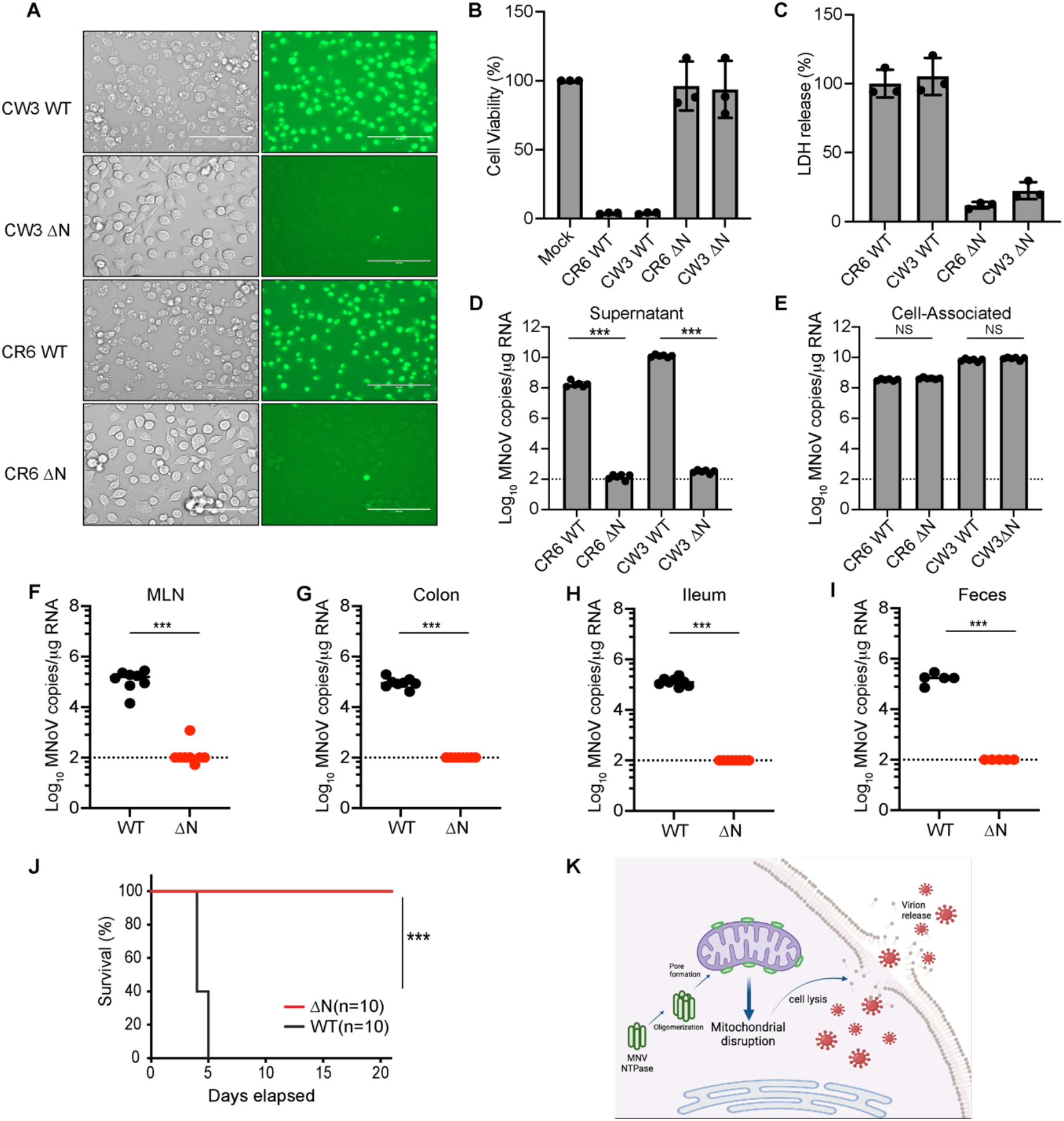
NTPase triggered cell death is essential for norovirus egress. (**A**) Representative images of Sytox-Green-stained BV2 cells infected with WT (MNoV^CR6^ and MNoV^CW3^) and mutant (MNoV^CR6^ΔN and MNoV^CW3^ΔN) MNoV at MOI=5 for 12 h. (**B**) Viability of BV2 cells infected with WT (MNoV^CR6^ and MNoV^CW3^) or mutant (MNoV^CR6^ΔN and MNoV^CW3^ΔN) MNoV at MOI=5 for 24 h. (**C**) LDH released from BV2 cells infected with WT (MNoV^CR6^ and MNoV^CW3^) and mutant (MNoV^CR6^ΔN and MNoV^CW3^ΔN) MNoV at MOI= 5 for 24 h. (**D-E**) BV2 cells infected with WT (MNoV^CR6^ and MNoV^CW3^) and mutant (MNoV^CR6^ΔN and MNoV^CW3^ΔN) MNoV at MOI=1 and viral genomes that were in the supernatant (**D**) or cell-associated (intracellular) (**E**) were quantified by qPCR at 24 h. (**F-J**) WT mice were challenged with 10^6^ PFU of MNoV^CR6^ and MNoV^CR6^ΔN perorally. Viral genomes were quantified by qPCR in the (**F**) MLN, (**G**) colon, (**H**) ileum, (**I**) feces at 7 days post infection. (**J**) Survival of *Stat1*^*− /−*^ mice after challenge with 10^6^ PFU of MNoV^CW3^ and MNoV^CW3^ΔN. Dashed line represents the limit of detection. (**K**) Model of NTPase triggered cell death for norovirus egress. Data represented in (**B-E)** are expressed as mean values ± s.d. from three technical replicates. Data are representative of three independent experiments. Statistical analysis for B-E was conducted using two-way ANOVA followed by Tukey’s multiple comparison test, For J, a log-rank (Mantel-Cox) test was performed. NS, not significant; *P<0.05; **P<0.01; ***P<0.001.

To determine whether NTPase-NT triggered cell death was essential for viral infection *in vivo*, we challenged mice with wildtype and mutant viruses. The CR6 strain of MNoV replicates in the intestines and is shed in the feces (*29*). We infected mice with MNoV^CR6^, MNoV^CR6^ΔN, or MNoV^CR6^ΔN20 and measured viral genomes in intestinal tissue and feces seven days post-infection (dpi). MNoV^CR6^ was detectable in the ileum, colon, MLN and feces. In contrast, viral genomes were rarely detected in intestinal tissues and feces of mice challenged with MNoV^CR6^ΔN and MNoV^CR6^ΔN20 (Fig. 4F-I and Fig. S21B-E). To further test whether NTPase N-terminal domain was essential for MNV *in vivo* replication, we orally inoculated *Stat1*^*-/-*^ mice with MNoV^CW3^, MNoV^CW3^ΔN, or MNoV^CW3^ΔN20. Consistent with prior studies, MNoV^CW3^ infection resulted in 100% lethality of *Stat1*^*-/-*^ mice (Fig. 4J) (*30*). However, we observed no lethality in *Stat1*^*-/-*^ mice infected with MNoV^CW3^ΔN and MNoV^CW3^ΔN20 viruses (Fig. 4J and Fig. S21F). These data suggest that NTPase-NT and mitochondrial localization of NTPase-NT is essential for MNoV replication *in vivo*.

In summary, norovirus NTPase-NT directly initiates programmed cell death by targeting mitochondrial outer membrane, leading to permeabilization of mitochondria (Fig. 4K). This mitochondrial permeabilization by NTPase is required for viral egress *in vitro* and replication *in vivo*. The N-terminal domain of NTPase mimics the cell death domain of MLKL, but rather than targeting the plasma membrane, NTPase targets mitochondria. Thus, the NTPase four helix bundle domain directly compromises mitochondrial function. These results suggest that mitochondrial integrity is a key checkpoint for not only apoptosis and pyroptosis, but also virus-induced programmed cell death (*31*–*33*). Sequence comparison of noroviruses and host MLKL protein sequences, suggests that *caliciviruses* may have stolen the four-helix bundle domain from a host. Our data using mutant viruses suggests that noroviruses co-opted this domain to induce cellular lysis for viral egress.

In the evolutionary arms race between viral fitness and host defense, viruses steal host proteins and repurpose them for viral replication. MLKL activity is targeted by many different viruses, including Poxviruses (*34, 35*). Interestingly, poxviruses encode a mimic derived from the regulatory domain of MLKL - the kinase-like domain – which suppresses necroptosis (Fig. 2I) (*33*–*36*). Noroviruses, on the other hand, have repurposed a MLKL-like four-helix bundle – the executioner domain - to facilitate virus replication. In summary, our study demonstrates a novel strategy for regulating programmed cell death and viral egress which may also be used by other viruses.

A family of viral proteins that cause membrane permeability are called viroporins (*39, 40*). These single or double transmembrane proteins homo-oligomerize and form ion channels in ER or plasma membranes. We propose that norovirus NTPases represent a novel pore forming protein, distinct from viroporins, based on the putative four-helix bundle structure of NTPase, homology to MLKL, and the specific localization of NTPase to the mitochondria (*36, 37*).

Our findings that noroviruses encode a protein to induce cell death challenges the widely held view that virus-triggered programmed cell death is a host survival strategy. While programmed cell death of host cells limits viral replication for many viruses, noroviruses actively induce cell death to facilitate viral spread. Given that NTPase is essential for virus genome replication and is a cell death executor during norovirus infection, it is possible that noroviruses regulate the timing, level of expression, and the localization of this multifunction protein. Because viral egress is a rate-limiting step for viral infection, our work illuminates a new putative target for therapeutics and vaccine design.

## Acknowledgments

We thank Z.G.Wang for providing pLVX-TRE3G vector and *Ripk3*^*− /−*^ mice. We thank Z.G.Wang and K.Yang for helpful discussions and technical assistance. We thank members of the Reese laboratory for technical assistance.

## Author contributions

G.X.W. and T.R. conceived the study. G.X.W. and D.Z. performed experiments and analyzed data.

D.H. performed sequence and phylogenetic analysis. R.C.O. provided reagents and expertise. G.X.W. and T.R. wrote the paper. All authors read and edited the manuscript.

## Data and materials availability

All data are available in the main paper and supplementary materials. All reagents are available from authors under a material transfer agreement with University of Texas Southwestern Medical Center.

