## supplemental figures for "Norovirus MLKL-like pore forming protein initiates programed cell death for viral egress"

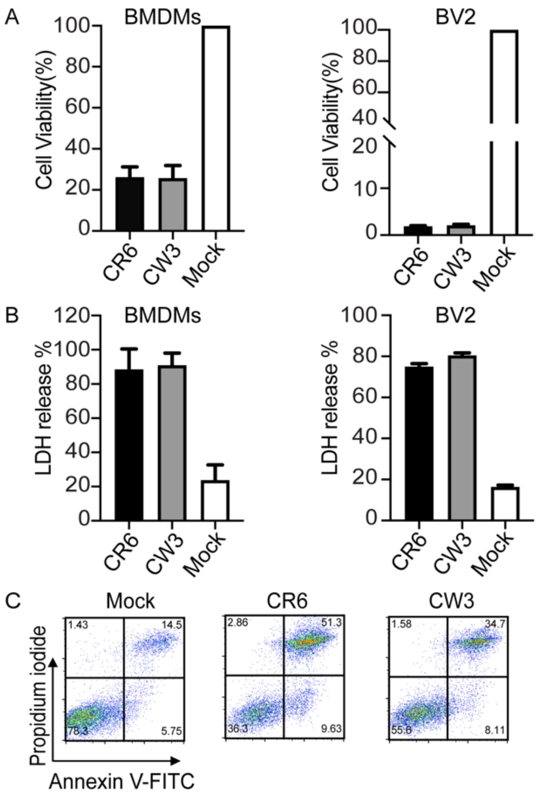


**Fig. S1. Noroviruses infection triggers programmed cell death.** (**A**) Viability of BMDMs and BV2 infected with $MNoV^{CW3}$and $MNoV^{CR6}$ at a MOI of 5 for 12 h. (**B**) LDH released from BMDMs and BV2 infected with $MNoV^{CW3}$and $MNoV^{CR6}$ at a MOI of 5 for 12 h. (**C**) Representative flow cytometry pseudocolor dot plots of propidium iodide (PI) and annexin V-stained BMDMs infected with $MNoV^{CW3}$ and $MNoV^{CR6}$ at a MOI of 5 for 12 h. Data (**A** and **B**) represented are expressed as mean values ± s.d. from three technical replicates. Data are representative of three independent experiments.

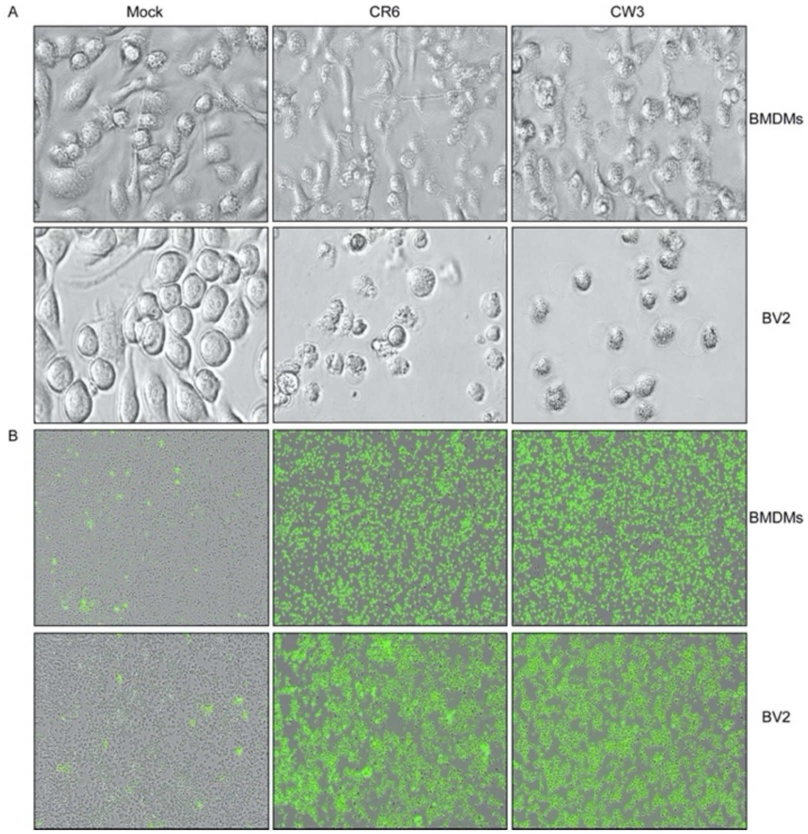


**Fig. S2**. **Noroviruses infection triggers programmed cell death.** (**A**) Bright-field images of BMDMs and BV2 infected with $MNoV^{CW3}$ and $MNoV^{CR6}$ at a MOI of 5 for 12 h. (**B**) Representative IncuCyte images of SytoxGreen-stained BMDMs and BV2 infected with $MNoV^{CW3}$ and $MNoV^{CR6}$ at a MOI of 5 for 24 h. All data shown are representative of at least three independent experiments.

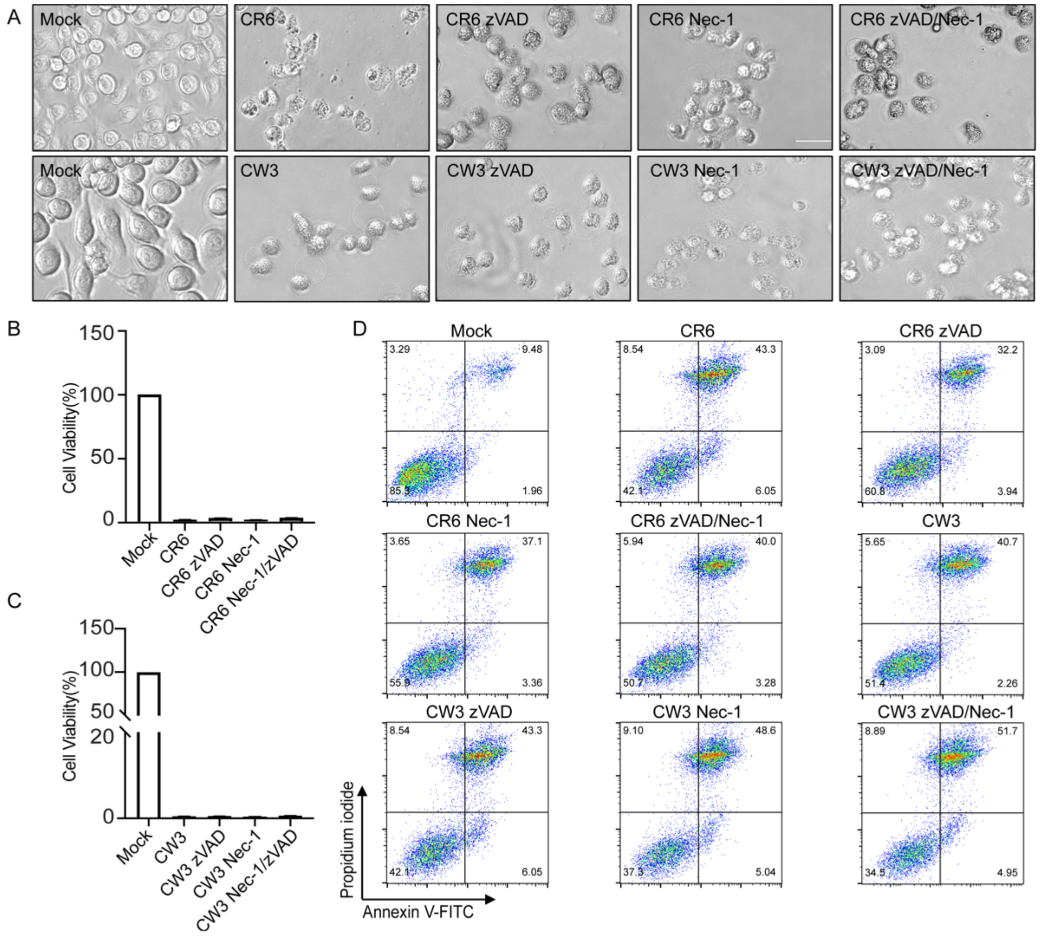


**Fig. S3. Caspase inhibitor and necroptosis inhibitor fail to block norovirus-induced cell death.** (**A**) Representative bright-field images of BMDMs infected with $MNoV^{CR6}$ or $MNoV^{CW3}$ at a MOI of 5 with or without zVAD and/or necrostatin-1 (Nec-1) for 12 h. (**B** to **C**) Cytotoxicity of BMDMs infected with (**B**) $MNoV^{CR6}$ or (**C**) $MNoV^{CW3}$ at a MOI of 5 with or without zVAD and/or Nec-1 for 24 h. (**D**) Representative flow cytometry pseudocolor dot plots of propidium iodide (PI) and annexin V-stained BMDMs are shown. BMDMs infected with $MNoV^{CW3}$ and $MNoV^{CR6}$ at a MOI of 5 with or without zVAD and/or Nec-1 for 12 h. All data shown are representative of at least three independent experiments. Data (**B** and **C**) represented are expressed as mean values ± s.d. from three technical replicates. Data are representative of three independent experiments.

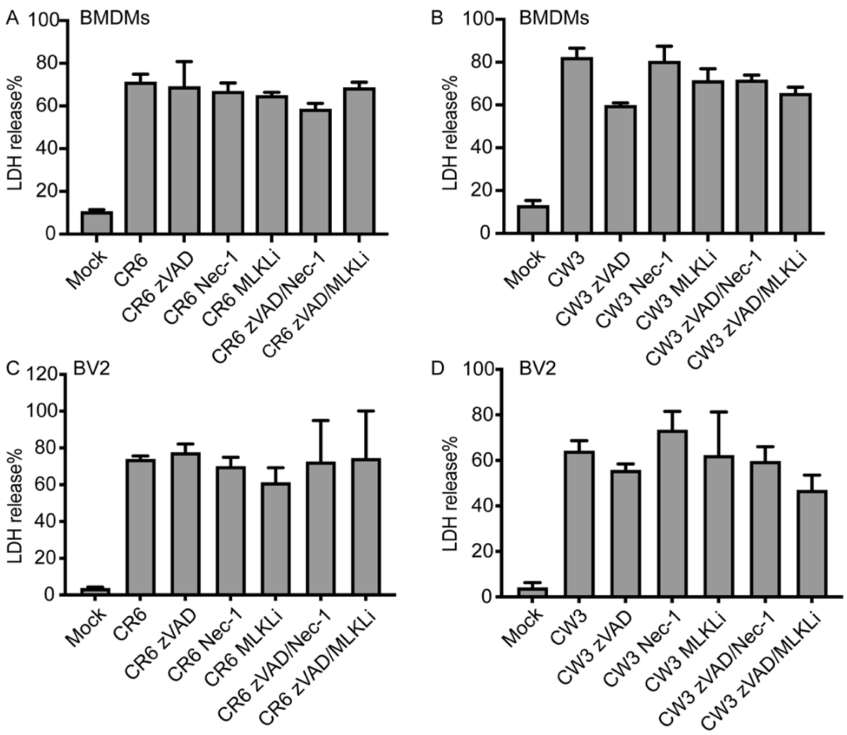


**Fig. S4. Pyroptosis, necroptosis, or apoptosis are not required for MNoV-induced cell death.** LDH released from BMDMs (**A** and **B**) and BV2 (**C** and **D**) infected with $MNoV^{CW3}$ and $MNoV^{CR6}$ (MOI=5) with or without zVAD, necrostatin-1 (Nec-1), MLKL inhibitor (MLKLi) either individually or in combination for 24 h. LDH release are expressed as mean ± s.d. from three technical replicates. Data are representative of three independent experiments.

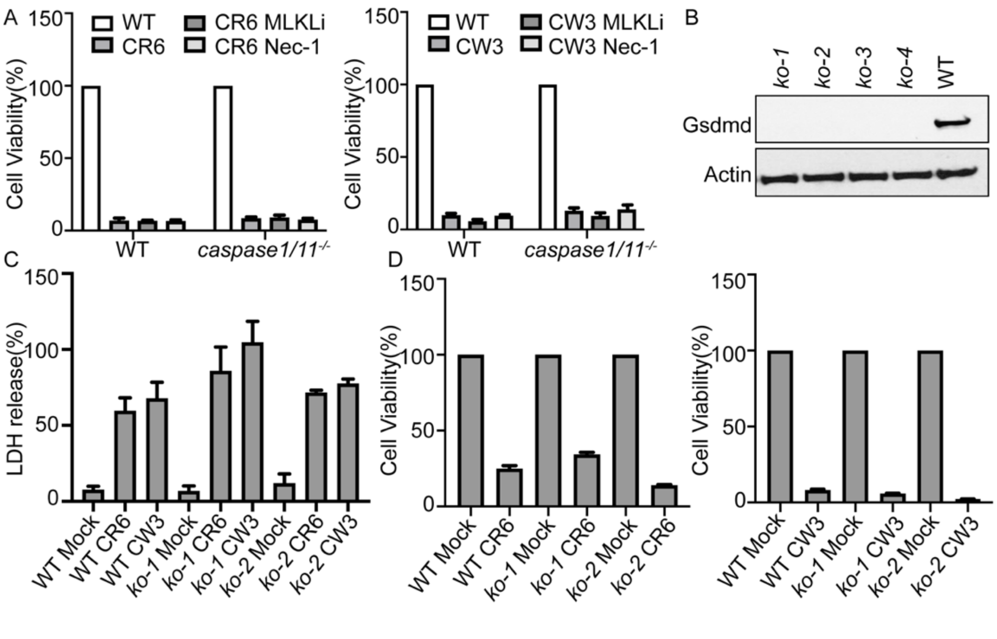


**Fig. S5. Pyroptosis is not required for MNoV-induced cell death.** (**A**) WT and *caspase1/11*$^{-/-}$ BMDMs cells were treated with vehicle, necrostatin-1 (Nec-1), or MLKL inhibitor (MLKLi) and infected with MNoV$^{CW3}$ and MNoV$^{CR6}$ at a MOI of 1 for 24 h. Cell viability measured by ATP levels. (**B**) GSDMD was deleted from BV2 cells by CRISPR genome editing. GSDMD expression levels in WT and 4 different clones of knockout BV2 cells. Lysates of indicated cells were blotted with anti-GSDMD antibody and anti-actin antibodies. (**C**) Comparison of LDH release in WT and *Gsdmd*$^{-/-}$ cells infected with MNoV$^{CW3}$and MNoV$^{CR6}$ (MOI=1) for 24 h. (**D**) Cell viability as measured by ATP levels WT and *Gsdmd*$^{-/-}$ cells infected with MNoV$^{CW3}$and MNoV$^{CR6}$ (MOI=1) for 24 h. Data (**A**, **C** and **D**) represented are expressed as mean values ± s.d. from three technical replicates. Data are representative of three independent experiments.

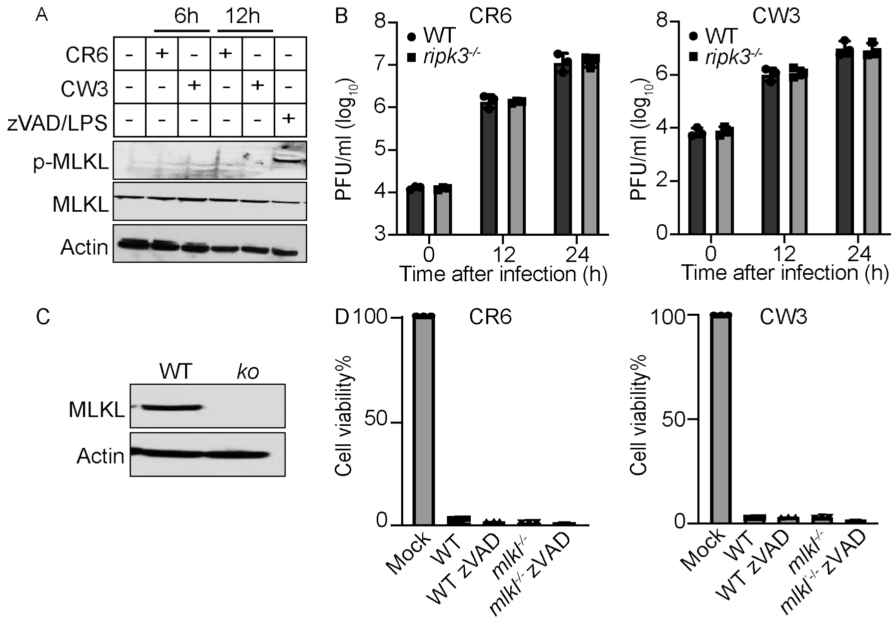


**Fig. S6. Necroptosis is not required for MNoV-induced cell death.** (**A**) WT BMDMs cells infected with $MNoV^{CW3}$ and $MNoV^{CR6}$ at a MOI of 1 or treated with 10 µM zVAD plus 100 ng/ml LPS. Lysates of indicated cells were blotted with anti-MLKL antibody, anti-pMLKL antibody and anti-actin antibodies. (**B**) WT and *Ripk3*$^{-/-}$ BMDMs cells infected with $MNoV^{CR6}$ or $MNoV^{CW3}$ at a MOI of 1 and virus was measured by plaque assay at indicated time. (**C**) MLKL was deleted from BV2 cells by CRISPR genome editing. Lysates of indicated cells were blotted with anti-MLKL antibody and anti-actin antibodies. (**D**) Comparison of ATP levels in $MNoV^{CW3}$and $MNoV^{CR6}$ (MOI=1) infected WT and *Mlkl*$^{-/-}$ cells with or without 10 µM zVAD for 24 h. Data (**D**) represented are expressed as mean values ± s.d. from three technical replicates. Data are representative of three independent experiments.

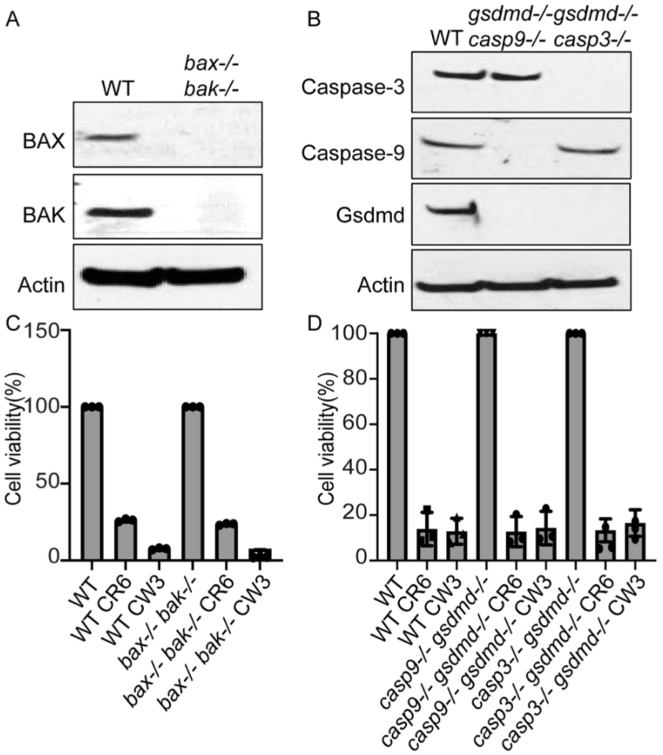


**Fig. S7. Apoptosis is not required for MNoV-induced cell death.** (**A**) *Bax* and *Bak* were deleted from BV2 cells by CRISPR genome editing. Lysates of WT or knockout cells were blotted with anti-Bak, anti-Bax and anti-actin antibodies. (**B**) *Gsdmd/Caspase-3* and *Gsdmd/Caspase-9* double knockout BV2 cells were made by CRISPR genome editing. Lysates of indicated cells were blotted with indicated antibodies. (**C**) Comparison of ATP levels in $MNoV^{CW3}$ and $MNoV^{CR6}$ (MOI=1) infected WT and $Bax^{-/-}$ / $Bak^{-/-}$ cells for 12 h. (**D**) Comparison of ATP levels in $MNoV^{CW3}$ and $MNoV^{CR6}$ (MOI=1) infected WT, $Gsdmd^{-/-}$ / $Caspase\text{-}3^{-/-}$ and $Gsdmd^{-/-}$ / $Caspase\text{-}9^{-/-}$ cells for 12 h. Data (**C** and **D**) represented are expressed as mean values ± s.d. from three technical replicates. Data are representative of three independent experiments.

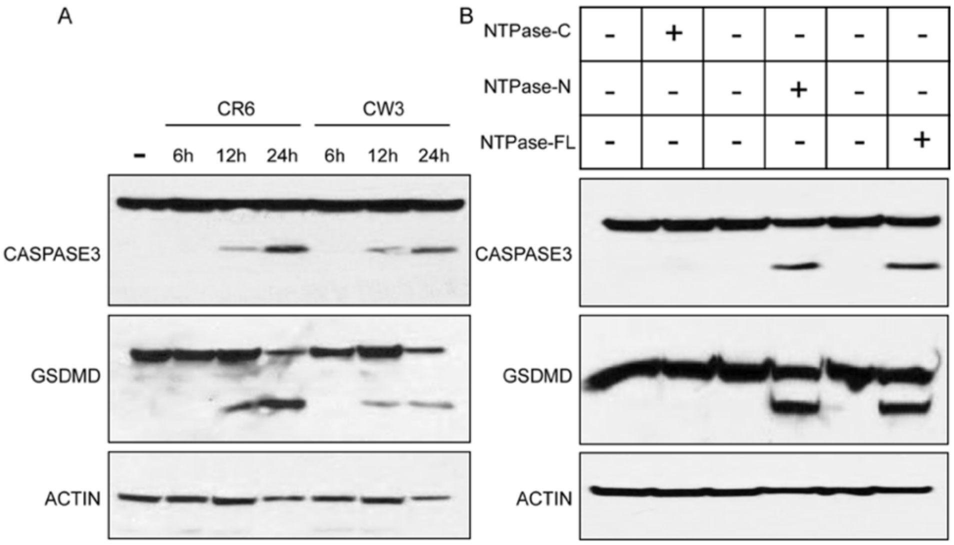


**Fig. S8. MNoV infection and expression Full-length, N-terminal NTPase of MNoV activated GSDMD and CASPASE-3 cleavage.** (**A**) Immunoblotting assays of norovirus infection-induced GSDMD and CASPASE-3 cleavage in BMDMs. BMDMs infected with MNoV$^{CW3}$and MNoV$^{CR6}$ (MOI=5) for 12h. (**B**) Immunoblotting assays of GSDMD and CASPASE-3 cleavage in BV2 cells stably expressing inducible Full-length, N- or C-terminal NTPase of MNoV strains CR6 in the absence or presence doxycycline for 12h. Results are representative of at least two independent experiments.

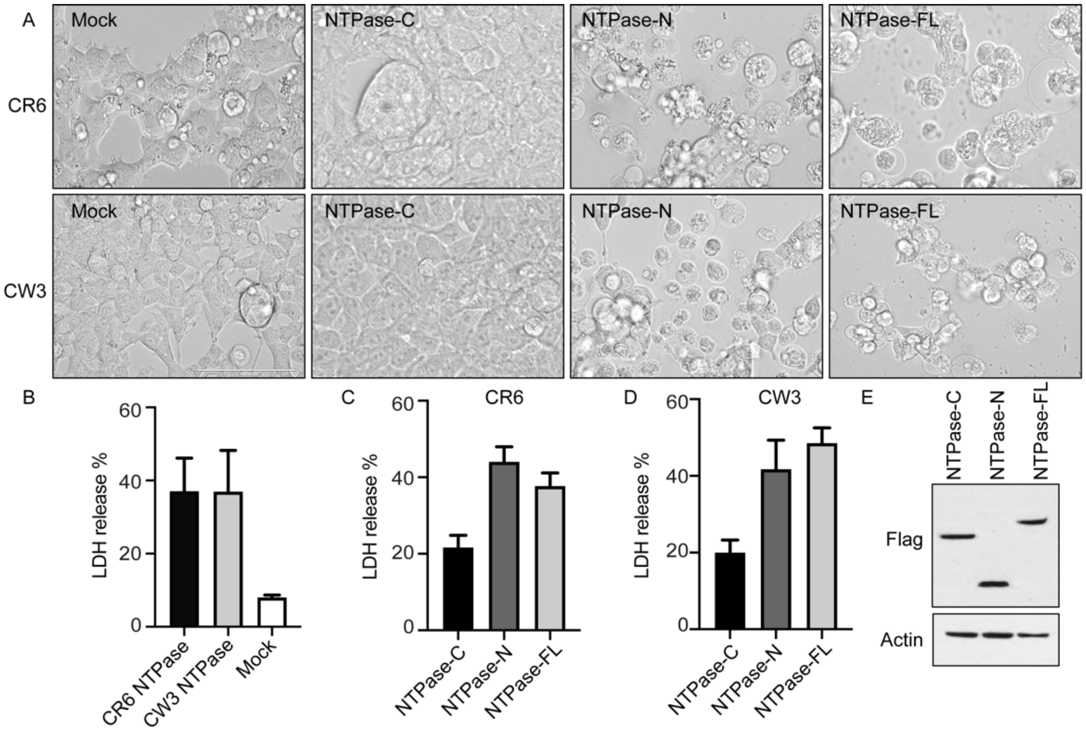


**Fig. S9. Norovirus NTPase directly triggers programmed cell death.** (**A**) Representative bright-field images of HEK293T cells transfected with Full-length, N- or C-terminal NTPase of MNoV strains CR6 or CW3. (**B**) LDH released from HEK293T cells transfected with Full-length NTPase of MNoV strains CR6 or CW3. (**C**) LDH released from HEK293T cells transfected with Full-length, N- or C-terminal NTPase of MNoV strains CR6. (**D**) LDH released from HEK293T cells transfected with Full-length, N- or C-terminal NTPase of MNoV strains CW3. LDH release is expressed as mean ± s.d. from three technical replicates. Data are representative of three independent experiments. (**E**) The immunoblot shows expression of HEK293T cells transfected with Full-length, N- or C-terminal of NTPase. All data shown are representative of three independent experiments.

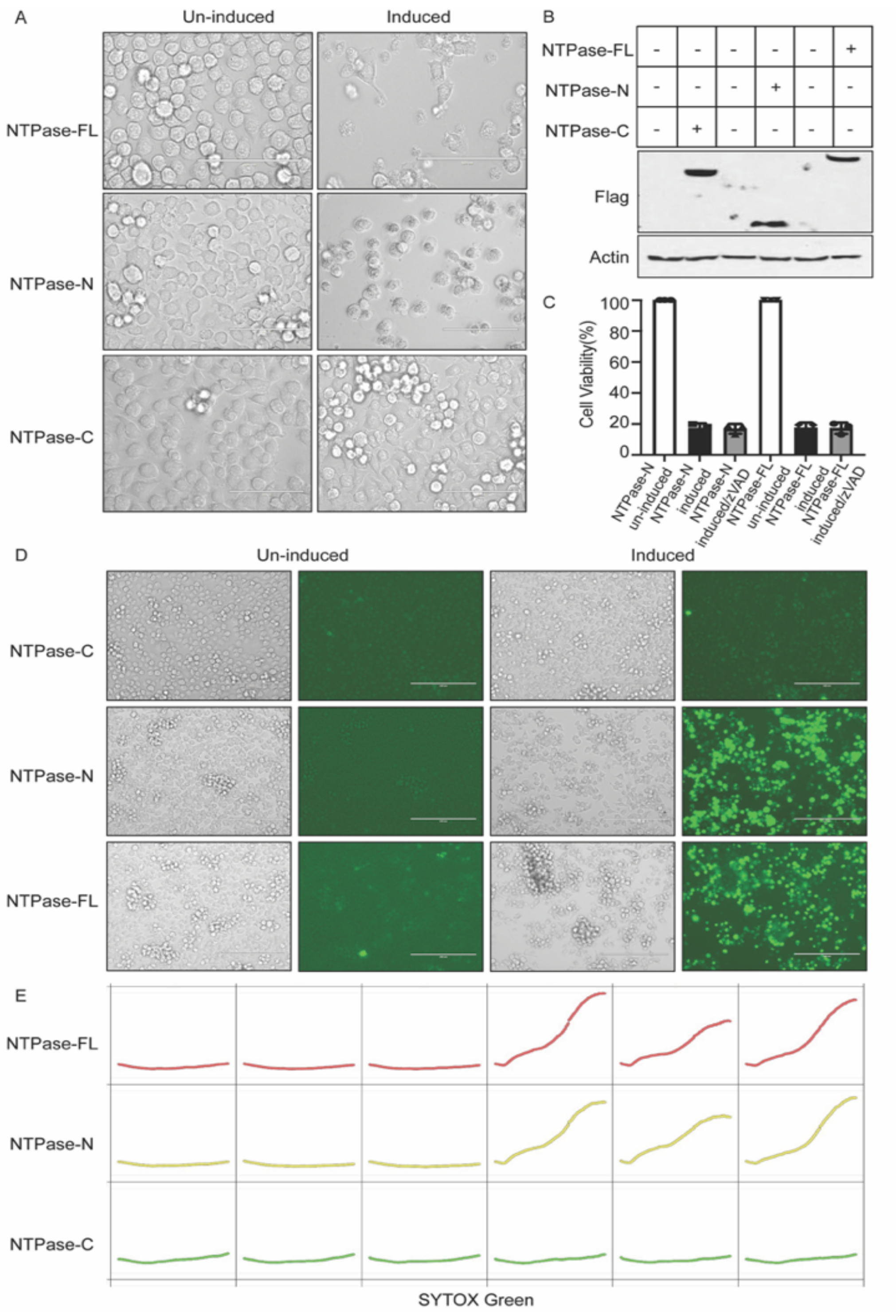
A
Un-induced
Induced
NTPase-FL
NTPase-N
NTPase-C
B
NTPase-FL
NTPase-N
NTPase-C
Flag
Actin
C
Cell Viability(%)
100
80
60
40
20
0
NTPase-N un-induced
NTPase-N induced
NTPase-N induced/zVAD
NTPase-FL un-induced
NTPase-FL induced
NTPase-FL induced/zVAD
D
Un-induced
Induced
NTPase-C
NTPase-N
NTPase-FL
E
NTPase-FL
NTPase-N
NTPase-C
SYTOX Green

**Fig. S10. Inducible expression NTPase of MNoV triggers programmed cell death**. Full-length, N- or C-terminal of NTPase fused C-terminal to Flag were stably expressed in BV2 cells under a tetracycline-inducible promoter. Doxycycline treatment to induced NTPase-FL, NTPase-N and NTPase-C expression. (**A**) Representative bright-field images of BV2 cells stably expressing inducible constructs in the absence or presence doxycycline for 12 h. (**B**) Immunoblot of NTPase with anti-Flag antibody in the absence or presence doxycycline for 12 h. (**C**) Cytotoxicity of BV2 cells expressing inducible constructs after doxycycline addition with or without 50 μM zVAD for 12 h. (**D**) Representative IncuCyte images of SytoxGreen-stained stably BV2 cells after doxycycline addition for 24 h and IncuCyte quantification over time. (**E**) Incucyte quantification, data are presented as mean of at least triplicate samples. Data represented are expressed as mean values ± s.d. from three technical replicates. Data are representative of three independent experiments.

A

B

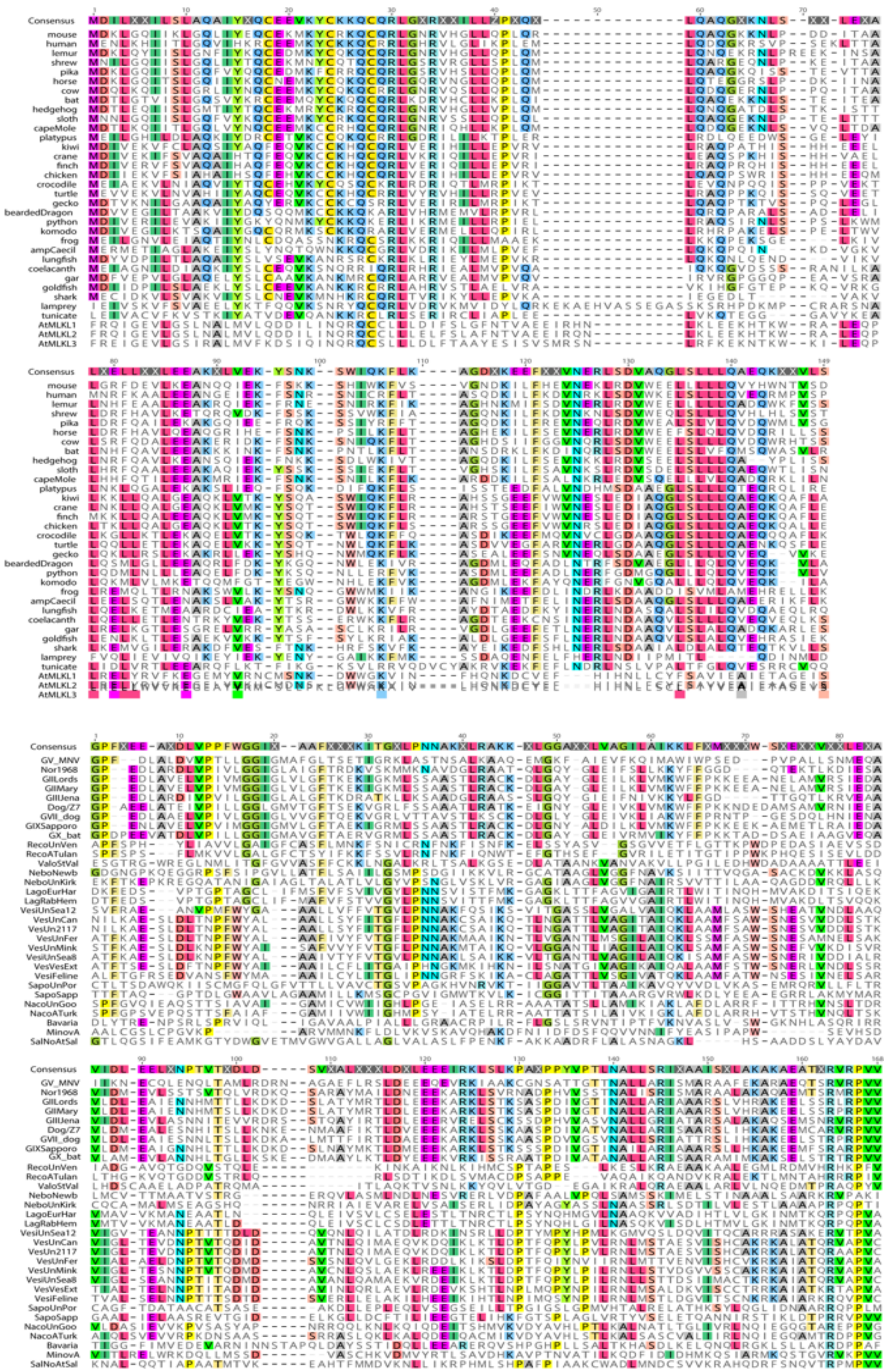
Consensus
mouse
human
lemur
shrew
pika
horse
cow
bat
hedgehog
sloth
capeMole
platypus
kiwi
crane
finch
chicken
crocodile
turtle
gecko
beardedDragon
python
komodo
frog
ampCaecil
lungfish
coelacanth
gar
goldfish
shark
lamprey
tunicate
AtMLKL1
AtMLKL2
AtMLKL3
GV_MNV
Nor1968
GIILords
GIIMary
GIIJena
Dog/Z7
GVII_dog
GIXSapporo
GX_bat
RecoUnVen
RecoATulan
ValoStVal
NeboNewb
NeboUnKirk
LagoEurHar
LagRabHem
VesiUnSea12
VesUnCan
VesUn2117
VesUnFer
VesUnMink
VesiUnSea8
VesVesExt
VesiFeline
SapoUnPor
SapoSapp
NacoUnGoo
NacoATurk
Bavaria
MinovA
SalNoAtSal

**Fig. S11. Expanded alignment of *calicivirus* 4HB domain sequences.** (**A**) Sequences were retrieved from NCBI. Viral species and strains were selected based on ICTV. Accession numbers are present in supplemental table 1. Sequences were aligned using MUSCLE in Geneious Prime. Residues matching consensus are colored. (**B**) NS3 N-terminus amino acid multiple sequence alignment (residues 1-158 relative to MNV) for thirty-one calicivirus species including representatives from norovirus genogroups. Sequences were retrieved from NCBI. Viral species and strains were selected based on ICTV. Accession numbers are present in supplemental table 1. Sequences were aligned using MUSCLE in Geneious Prime. Residues matching consensus are colored.

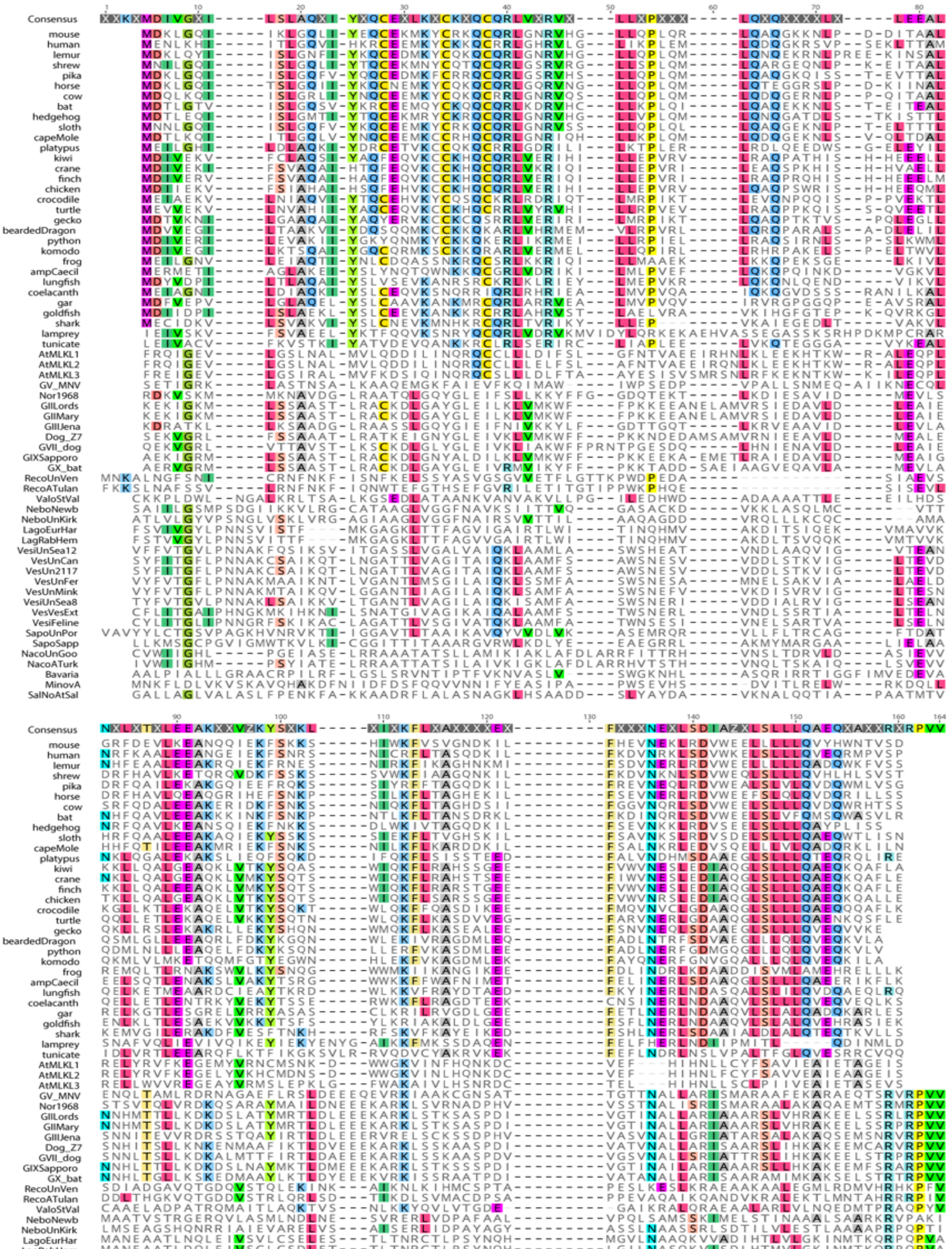
Consensus
mouse
human
lemur
shrew
pika
horse
cow
bat
hedgehog
sloth
capeMole
platypus
kiwi
crane
finch
chicken
crocodile
turtle
gecko
beardedDragon
python
komodo
frog
ampCaecil
lungfish
coelacanth
gar
goldfish
shark
lamprey
tunicate
AtMLKL1
AtMLKL2
AtMLKL3
GV_MNV
Nor1968
GIILords
GIIMary
GIIIJena
Dog_Z7
GVII_dog
GIXSapporo
GX_bat
RecoUnVen
RecoATulan
ValoStVal
NeboNewb
NeboUnKirk
LagoEurHar
LagRabHem
VesiUnSea12
VesUnCan
VesUn2117
VesUnFer
VesUnMink
VesiUnSea8
VesVesExt
VesiFeline
SapoUnPor
SapoSapp
NacoUnGoo
NacoATurk
Bavaria
MinovA
SalNoAtSal

**Fig. S12. Expanded alignment of cellular and viral 4HB domain sequences.** Sequences were retrieved from NCBI. Viral species and strains were selected based on ICTV. Accession numbers are present in supplemental table 1. Sequences were aligned using MUSCLE in Geneious Prime. Residues matching consensus are colored.

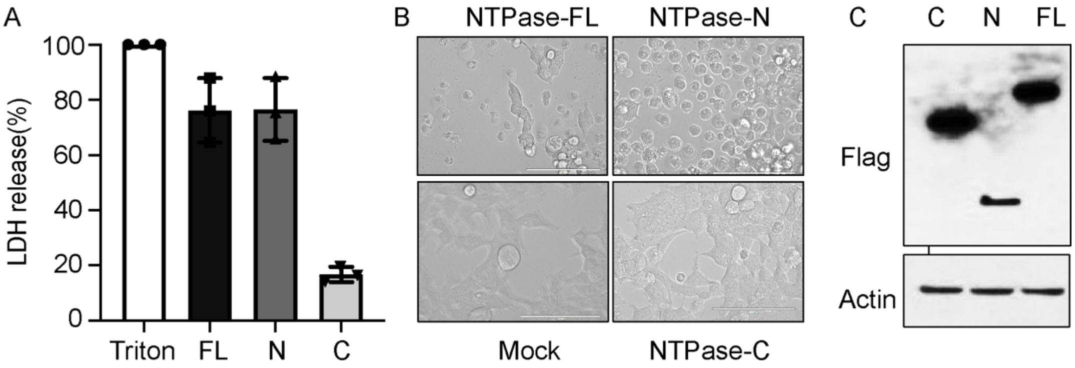


**Fig. S13. HNoV NTPase directly triggers programmed cell death.** (**A**) LDH released from HEK293T cells transfected with Full-length, N- or C-terminal NTPase of HNoV MD145. (**B**) Representative bright-field images of HEK293T cells transfected with Full-length, N- or C-terminal NTPase of MD145. (**C**) The immunoblot shows expression of transfected Full-length, N- or C-terminal NTPase of MD145. All data shown are representative of at least three independent experiments. Data (**A**) represented are expressed as mean values ± s.d. from three technical replicates. Data are representative of three independent experiments.

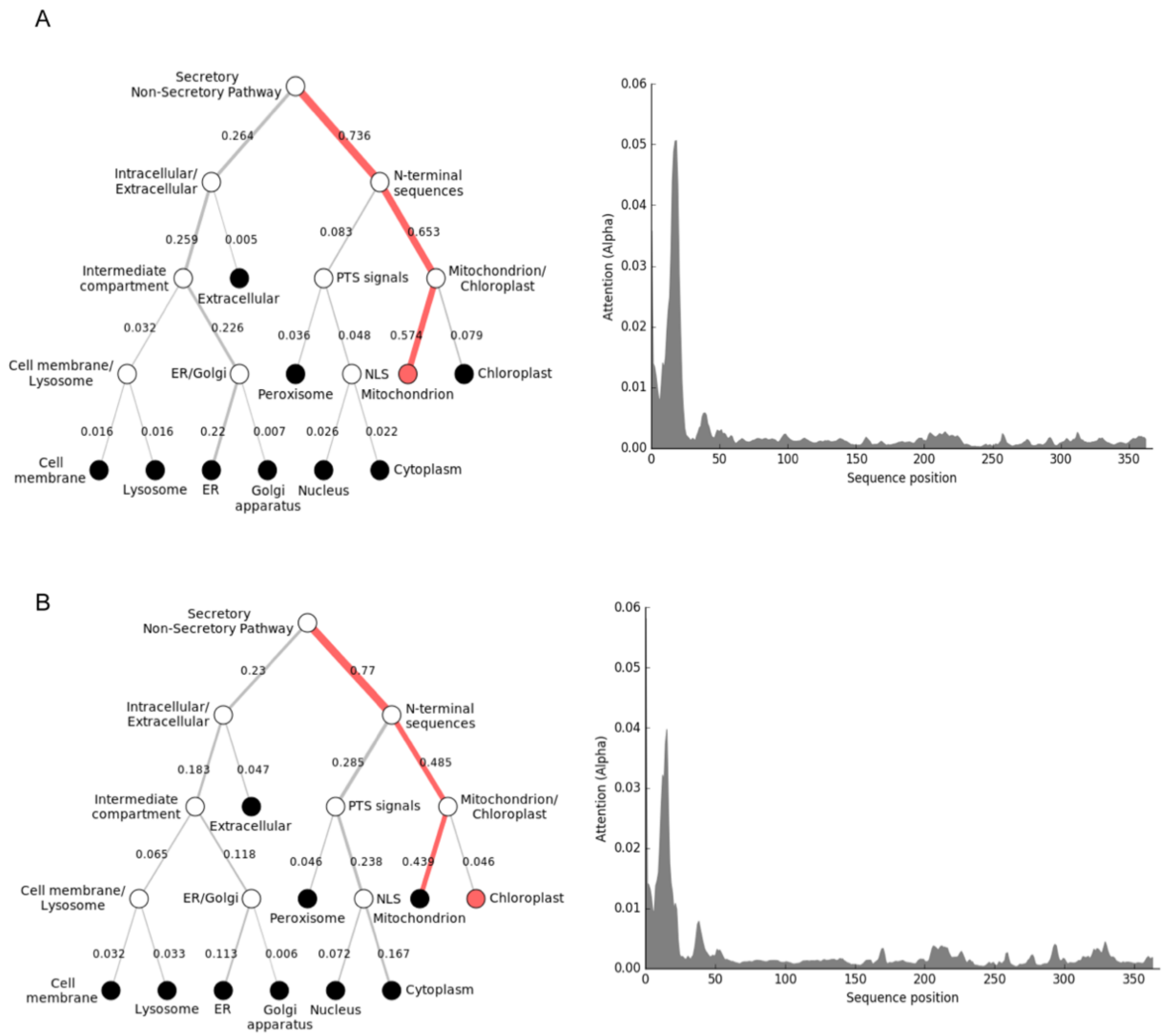


**Fig. S14. Subcellular localization prediction of NTPase.** DeepLoc prediction server revealed that full-length NTPase of (**A**) HNoV and (**B**) MNoV have high probability for mitochondrial localization.

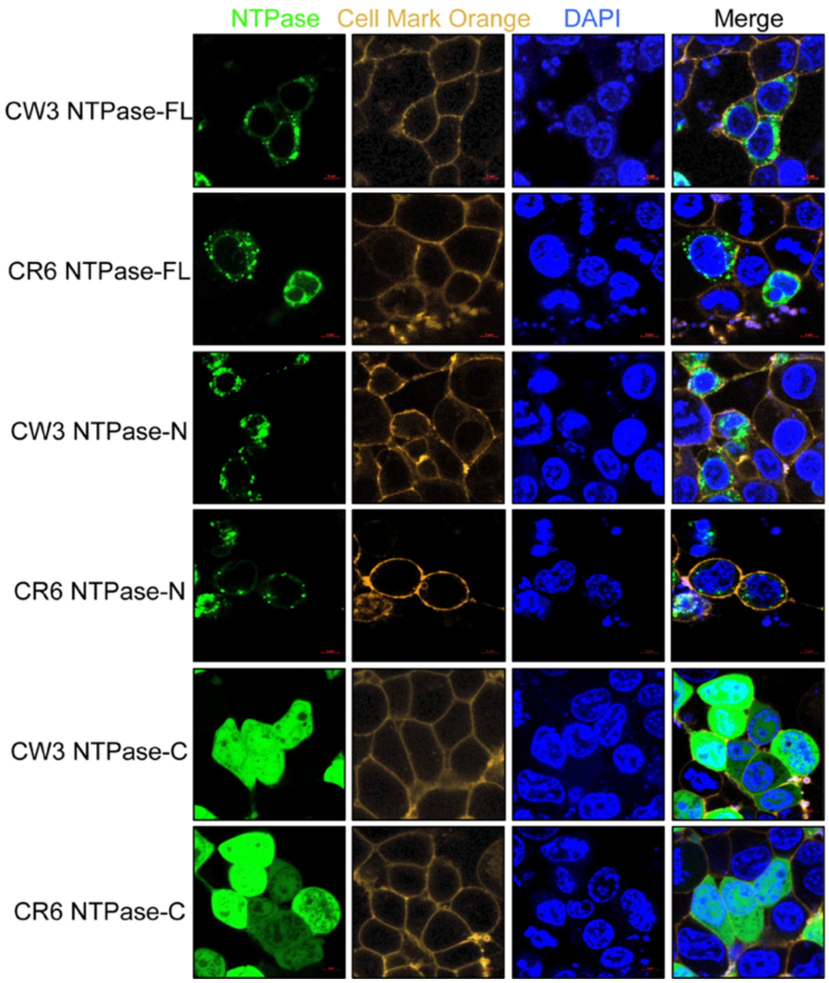


**Fig. S15. Analysis subcellular localization of NTPase.** Representative microscopy imaging of NTPase subcellular localization in HEK293T cells. Flag-tagged NTPase-FL, NTPase-N and NTPase-C transfected HEK293T cells, the distribution of NTPase (green) was detected by immunofluorescence with cellular membrane marker (yellow). Scale bars, 10 μm. Results are representative of at least two independent experiments.

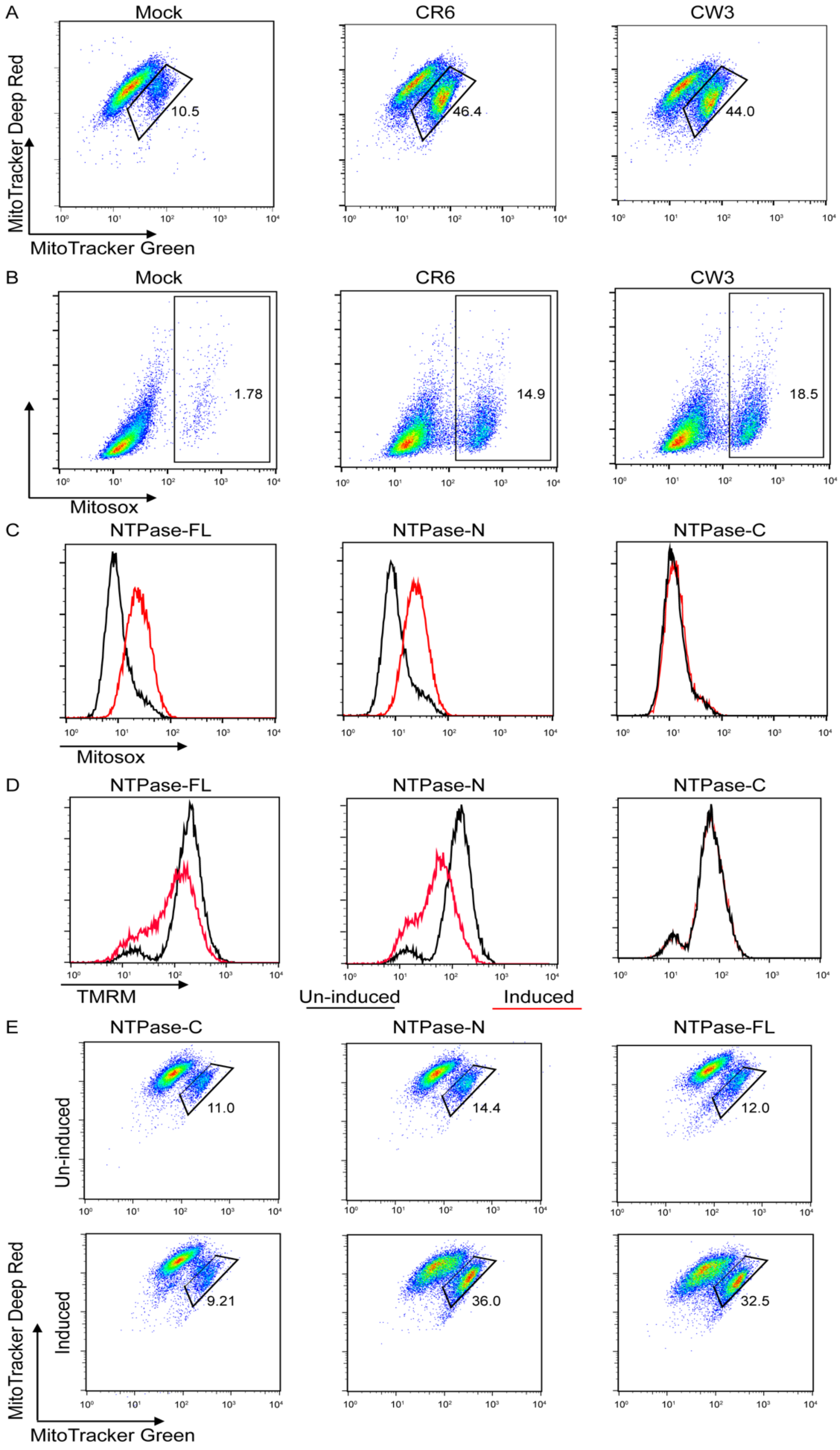
A
Mock
CR6
CW3
MitoTracker Deep Red
MitoTracker Green
10.5
46.4
44.0
B
Mock
CR6
CW3
Mitosox
1.78
14.9
18.5
C
NTPase-FL
NTPase-N
NTPase-C
Mitosox
D
NTPase-FL
NTPase-N
NTPase-C
TMRM
Un-induced
Induced
E
NTPase-C
NTPase-N
NTPase-FL
Un-induced
Induced
11.0
14.4
12.0
9.21
36.0
32.5
MitoTracker Deep Red
MitoTracker Green

**Fig. S16. MNoV infection and expression Full-length, N-terminal NTPase of MNoV altered mitochondria membrane potential and increased mitochondrial ROS abundance.** (**A**) Flow cytometric analysis of mitochondrial status in macrophages infected with MNoV$^{CW3}$and MNoV$^{CR6}$ at a MOI of 5 for 12 h. Gates represent cells with damaged mitochondria. (**B**) Flow cytometric analysis of mitochondrial ROS in macrophages infected with MNoV$^{CW3}$and MNoV$^{CR6}$ at a MOI of 5 for 12 h. (**C-E**) Full-length, N- or C-terminal of NTPase fused to a C-terminal Flag tag were stably expressed in BV2 cells under a tetracycline-inducible promoter. Doxycycline was used to induce NTPase-FL, NTPase-N and NTPase-C expression. (**C**) Flow cytometric analysis of mitochondrial ROS in the absence (black) or presence (red) of doxycycline. (**D**) Flow cytometric analysis mitochondrial membrane potential (Ψm) of stably BV2 cells in the absence (black) or presence (red) of doxycycline measured by TMRM fluorescence. (**E**) Flow cytometric analysis of mitochondrial status in the absence or presence doxycycline. Gates represent cells with damaged mitochondria. Experiments were performed at least two independent times each in triplicate.

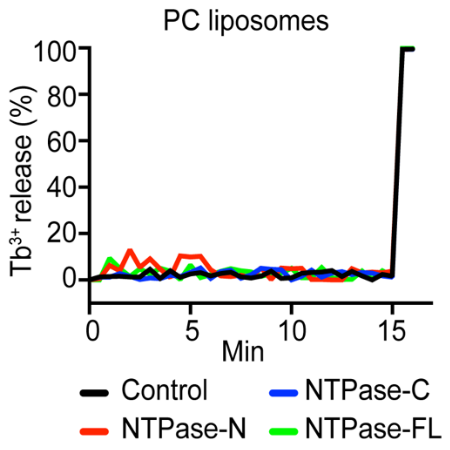


**Fig. S17**. **NTPase-FL and NTPase-N did not lyse PC liposomes.** Purified Full-length, N- or C-terminal NTPase proteins were incubated with liposomes. Liposome leakage was monitored by measuring DPA chelating-induced fluorescence of released $Tb^{3+}$ relative to that of Triton X-100 treatment.

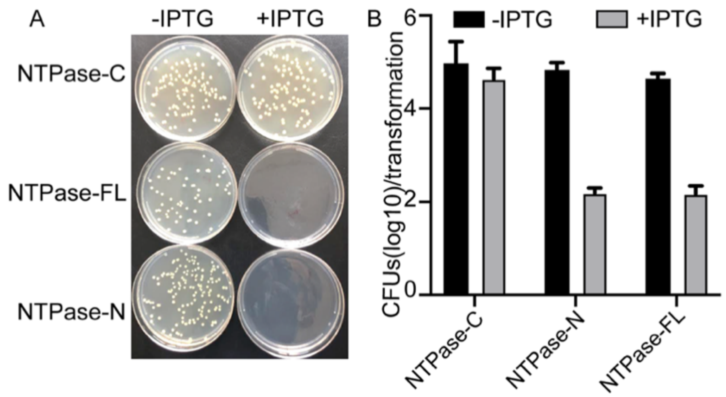


**Fig. S18. NTPase-FL and NTPase-N are toxic to *Escherichia coli.*** (**A**) Representative agar plates showing transformed *E. coli* colonies for NTPase-FL, NTPase-N and NTPase-C. (**B**) Bacterial colony-forming units (CFU) per transformation for NTPase-FL, NTPase-N and NTPase-C are shown in the logarithmic form (log10) from three technical replicates.

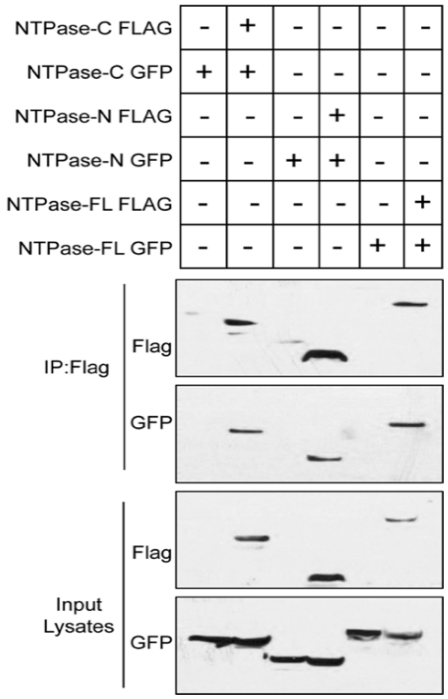


**Fig. S19. Self-associated of Full-length, N- and C-terminal NTPase of MNoV.** Flag immunoprecipitation of lysates of HEK293T cells, transfected with GFP-NTPase-FL, GFP-NTPase-N, GFP-NTPase-C and/or Flag-NTPase-FL, Flag-NTPase-N, Flag-NTPase-C, were analyzed by immunoblot with indicated antibodies.

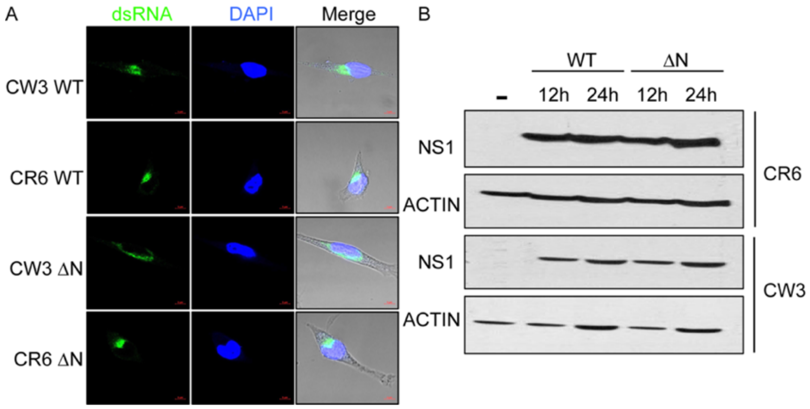


**Fig. S20. Mutant MNoV lacking NTPase N-terminus form replication complexes and normally express viral non-structural proteins.** (**A**) Representative confocal microscopy images dsRNA (green) in BMDMs infected with WT ($MNoV^{CW3}$and $MNoV^{CR6}$) or mutant ($MNoV^{CW3}\Delta N$ and $MNoV^{CR6}\Delta N$) virus at a MOI of 5 for 12 h. (**B**) BMDMs derived from wild-type mice were infected with WT ($MNoV^{CW3}$and $MNoV^{CR6}$) or mutant ($MNoV^{CW3}\Delta N$ and $MNoV^{CR6}\Delta N$) viruses. Cells collected at indicated time points and total cell lysates were subjected to anti-NS1 or anti-actin immunoblotting.

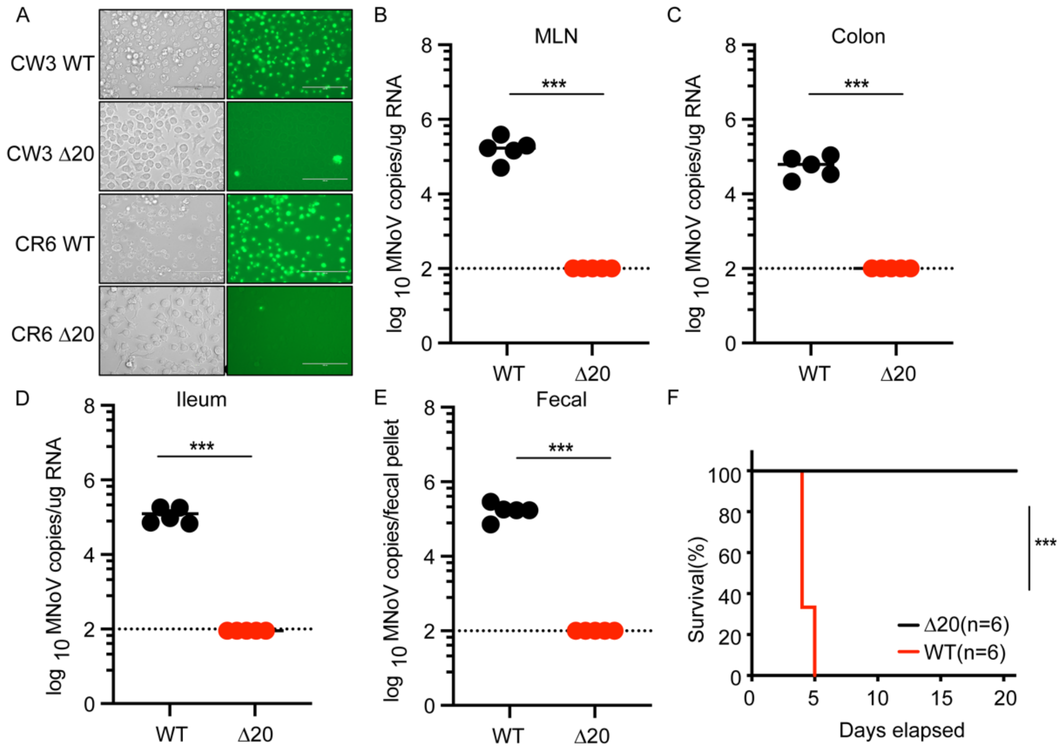


**Fig. S21. Mitochondrial localization of NTPase is essential for inducing cell death and norovirus egress.** (**A**) Representative images of Sytox-Green-stained BV2 cells infected with WT ($MNoV^{CR6}$ and $MNoV^{CW3}$) and mutant ($MNoV^{CR6}$ΔN20 and $MNoV^{CW3}$ΔN20). WT mice were challenged with $10^6$ PFU of $MNoV^{CR6}$ and $MNoV^{CR6}$ΔN20 perorally. All mice survived infection. Viral genomes were quantified in the (**B**) MLN, (**C**) colon, (**D**) ileum, (**E**) feces at 7 days post infection. (**F**) Survival of *Stat1*$^{-/-}$ mice after challenge with $10^6$ PFU of $MNoV^{CW3}$ and $MNoV^{CW3}$ΔN20. Dashed line represents the limit of detection. Statistical analysis in B through E was conducted using two-way ANOVA followed by Tukey's multiple comparison test. P-value for F was calculated with a log-rank (Mantel-Cox) test, NS, not significant; *P<0.05; **P<0.01; ***P<0.001.

Table 1: Calicivirus accession numbers for sequences used in this study

| | Genus | Species | Virus name | Isolate | Virus abbreviation | Alignment label in supplemental data | Genome accession | ORF1 protein accession |
|---|---|---|---|---|---|---|---|---|
| 1 | *Bavovirus* | *Bavaria virus* | chicken calicivirus | V0021/DE/2004 | CCV | Bavaria | HQ010042 | ADN88287 |
| 2 | *Lagovirus* | *European brown hare syndrome virus* | | GD/FR/1989 | EBHSV | LagoEurHar | Z69620 | CAA93445.1 |
| 3 | *Lagovirus* | *Rabbit hemorrhagic disease virus* | rabbit hemorrhagic disease virus | GH/DE/1988 | RHDV | LagRabHem | M67473 | AAA47285.1 |
| 4 | *Minovirus* | *Minovirus A* | fathead minnow calicivirus | US/2012 | FMCV | MinovaA | KX371097 | AQM56929.1 |
| 5 | *Nacovirus* | *Nacovirus A* | turkey calicivirus | L11043/DE/2011 | TCV | NacoATurk | JQ347522 | AFH89833.1 |
| 6 | *Nacovirus* | *unclassified* | goose calicivirus strain N | | | NacoUnGoo | KJ473715 | AHY39267.1 |
| 7 | *Nebovirus* | *Newbury 1 virus* | Newbury-1 virus | UK/1976 | N1V | NeboNewb | DQ013304 | AAY60849.1 |
| 8 | *Nebovirus* | *unclassified* | Kırklareli virus | | | NeboUnKirk | KT119483 | ALC74362.1 |
| 9 | *Norovirus* | *Norwalk virus* | Norwalk virus | US/1968 | NV | Nor1968 | M87661 | AAB50465.1 |
| 10 | *Recovirus* | *Recovirus A* | Tulane virus | US/2004 | TV | RecoATulan | EU391643 | ACB38131.1 |
| 11 | *Recovirus* | *unclassified* | human recovirus Venezuela | | | RecoUnVen | MG571787 | AWU65998.1 |
| 12 | *Salovirus* | *Nordland virus* | Atlantic salmon calicivirus | Nordland/NO/2011 | ASCV | SalNoAtSal | KJ577139 | AHX24375 |
| 13 | *Sapovirus* | *Sapporo virus* | Sapporo virus | JP/1982 | SV | SapoSapp | HM002617 | ADG03646.1 |
| 14 | *Sapovirus* | *unclassified* | porcine_sapovirus_JJ681 | | | SapoUnPor | AY974192 | AAY40312.2 |
| 15 | *Valovirus* | *Saint Valerian virus* | St-Valérien calicivirus | AB90/CAN/2006 | SVCV | ValoStVal | FJ355928 | ACQ44559.1 |
| 16 | *Vesivirus* | *Feline calicivirus* | feline calicivirus | F9/US/1958 | FCV | VesiFeline | M86379 | AAA79326.1 |
| 17 | *Vesivirus* | *Vesicular exanthema of swine virus* | vesicular exanthema of swine virus | A48/US/1948 | VESV | VesVesExt | U76874 | AAG13641.1 |
| 18 | *Vesivirus* | *unclassified* | calicivirus 2117 | | 2117 | VesUn2117 | AY343325 | AAQ24845.2 |
| 19 | *Vesivirus* | *unclassified* | canine vesivirus [Bari/212/07/ITA] | | CaVV | VesUnCan | JN204722 | AFI58320.1 |
| 20 | *Vesivirus* | *unclassified* | mink calicivirus [MCV-DL/2007/CN] | | MCV | VesUnMink | JX847605 | AFY07854.1 |
| 21 | *Vesivirus* | *unclassified* | San Miguel sea lion virus 8 [274] | | SMSV-8 | VesiUnSea8 | KM244552; [illegible] | AIU47325.1 |
| 22 | *Vesivirus* | *unclassified* | San Miguel sea lion virus 12 [2615T] | | SMSV-12 | VesiUnSea12 | MK962352 | QCZ35343.1 |
| 23 | *Vesivirus* | *unclassified* | ferret-badger vesivirus [JX12/China/2012] | | FBV | VesUnFer | KJ701554 | AJO15926.1 |

Virus information was retrieved from ICTV (https://talk.ictvonline.org/ictv-reports/ictv_online_report/positive-sense-rna-viruses/w/caliciviridae). Sequences were retrieved from the NCBI database. Genome accessions linked to predicted ORF1 protein sequences, which were used in the analysis.

Table 2: Cellular MLKL accession numbers for animal and plant sequences used in this study

| | Species | Common name | MLKL protein accession |
|---|---|---|---|
| 1 | *Homo sapiens* | human | NP_689862.1 |
| 2 | *Galeopterus variegatus* | Sunda flying lemur | XP_008570998.1 |
| 3 | *Tupaia chinensis* | tree shrew | XP_006152702.1 |
| 4 | *Ochotona princeps* | pika | XP_004584250 |
| 5 | *Mus musculus* | mouse | NP_083281.1 |
| 6 | *Equus caballus* | horse | XP_005608486.1 |
| 7 | *Bos taurus* | cow | XP_002694753.1 |
| 8 | *Pteropus vampyrus* | bat | XP_023378937.1 |
| 9 | *Erinaceus europaeus* | West European hedgehog | XP_007527656.1 |
| 10 | *Choloepus didactylus* | southern two-toed sloth | XP_037672089.1 |
| 11 | *Chrysochloris asiatica* | Cape golden mole | XP_006860370.1 |
| 12 | *Ornithorhynchus anatinus* | platypus | XP_007655108.1 |
| 13 | *Apteryx rowi* | Okarito brown kiwi | XP_025944378.1 |
| 14 | *Balearica regulorum gibbericeps* | East African grey crowned-crane | XP_010297767.1 |
| 15 | *Geospiza fortis* | medium ground-finch | XP_005422873.1 |
| 16 | *Gallus gallus* | chicken | XP_015134716.1 |
| 17 | *Crocodylus porosus* | Australian saltwater crocodile | XP_019391158.1 |
| 18 | *Chelonia mydas* | green sea turtle | XP_007071298.3 |
| 19 | *Gekko japonicus* | gecko | XP_015285205.1 |
| 20 | *Pogona vitticeps* | central bearded dragon | XP_020653567.1 |
| 21 | *Python bivittatus* | Burmese python | XP_025027125.1 |
| 22 | *Varanus komodoensis* | Komodo dragon | XP_044290260.1 |
| 23 | *Xenopus tropicalis* | tropical clawed frog | XP_004914113.2 |
| 24 | *Microcaecilia unicolor* | caecilians | XP_030059489.1 |
| 25 | *Protopterus annectens* | West African lungfish | XP_043937895.1 |
| 26 | *Latimeria chalumnae* | coelacanth | XP_006010420.1 |
| 27 | *Lepisosteus oculatus* | gar | XP_015223778.1 |
| 28 | *Carassius auratus* | goldfish | XP_026058901.1 |
| 29 | *Carcharodon carcharias* | great white shark | XP_041046768.1 |
| 30 | *Petromyzon marinus* | lamprey | XP_032816483.1 |
| 31 | *Ciona intestinalis* | vase tunicate | XP_002122121.1 |
| 32 | *Arabidopsis thaliana* (AtMLK1) | thale cress | Mahdi *et al*. Cell Host Microbe 2020 |
| 33 | *Arabidopsis thaliana* (AtMLK2) | thale cress | Mahdi *et al*. Cell Host Microbe 2020 |
| 34 | *Arabidopsis thaliana* (AtMLK3) | thale cress | Mahdi *et al*. Cell Host Microbe 2020 |

Table 3. Norovirus accession numbers for sequences used in this study

| | Genus | Species | Virus name | Isolate | Virus abbreviation | Alignment label in supplemental data | Genome accession | ORF1 protein accession |
|---|---|---|---|---|---|---|---|---|
| 1 | *Norovirus* | *Norwalk virus* | Norwalk virus | US/1968 | NV | Nor1968 | M87661 | AAB50465.1 |
| 2 | *Norovirus* | *Norwalk virus* | Lordsdale virus | GII/UK/1995 | LV | GIILords | X86557 | CAA60254.1 |
| 3 | *Norovirus* | *Norwalk virus* | Maryland virus | GII/US/1987 | MV | GIIMary | AY032605 | AAK50354.1 |
| 4 | *Norovirus* | *Norwalk virus* | Jena virus | GIII/DE/1980 | JV | GIIIJena | AJ011099 | CAA09480.1 |
| 5 | *Norovirus* | *Norwalk virus* | murine norovirus 1 | GV/US/2002 | MNV1 | GV_MNV | AY228235 | AAO63098.2 |
| 6 | *Norovirus* | *Norwalk virus* | | GVI isolate Dog/Z7/19/CH | | Dog/Z7 | MW662289.1 | QSD75797.1 |
| 7 | *Norovirus* | *Norwalk virus* | dog norovirus | GVII/HK/2007 | CaV | GVII_dog | FJ692500 | ACV89839.1 |
| 8 | *Norovirus* | *Norwalk virus* | Sapporo-HK299 virus | GIX/JP/2007 | SaV | GIXSapporo | KJ196290 | AII73758.1 |
| 9 | *Norovirus* | *Norwalk virus* | bat norovirus | GX/CN/2010 | BtRsV | GX_bat | KJ790198 | AID69173.1 |

### Materials and Methods

Plasmids, antibodies and reagents

Nucleotide sequences of Norovirus genes were amplified with Q5 High-Fidelity 2 × Master Mix (NEB) from pCR6, pCW3 and pSPORT MD145 plasmids. Flag-tagged constructs of Norovirus genes were generated by cloning the indicated genes into pCMV-6b-Flag vector with N-terminal 3 × Flag tag using Gibson Assembly Master Mix (NEB). GFP-tagged constructs were assembled by subcloning the indicated genes into pEGFP-C2 vector for transient expression in HEK293T cells. Truncation mutants of the NTPase were constructed by the standard PCR cloning strategy and inserted into pEGFP-C2 vector and pCMV-6b-Flag vector. For recombinant expression in *E. coli*, the cDNAs were cloned into a pMAL-c6T × $His_6$-MBP vector (NEB). For inducible Tet-on overexpression, coding sequences of NTPase was amplified from pCR6 plasmids and subcloned into pLVX-TRE3G vector (Clontech) using Gibson Assembly Master Mix (NEB). All plasmids were verified by DNA sequencing.

Natural and synthetic lipid products used for liposome preparation were obtained from Avanti Polar Lipids Inc. Lipid strips were purchased from Echelon Biosciences Inc. MitoTracker Deep Red, MitoTracker Green, MitoSOX and TMRM were obtained from Invitrogen. Ultrapure LPS were obtained from InvivoGen. Doxycycline (D9891), terbium chloride ($TbCl_3$), polybrene (H9268) and DPA (dipicolinic acid) were purchased from Sigma-Aldrich. z-VAD (FMK007, R&D) was dissolved in DMSO. MLKL inhibitor was kindly provided by Z.G.Wang (UT Southwestern Medical Center). The antibodies used in this study include: anti-Flag (A8592, Sigma), anti-Actin (A2066, Sigma), anti-GFP (SC9996, Santa Cruz), anti-Cytochrome *C* (556433, BD Biosciences), anti-BAK (12105, Cell Signaling), anti-BAX (2772s, Cell Signaling), anti-MBP (E8032S, NEB), anti-MLKL (ZRB1142, Sigma), anti-pMLKL (196436, Abcam), anti-Caspase 3 (9962, Cell Signaling), anti-Caspase 9 (9504, Cell Signaling), anti-COX IV (4844, Cell Signaling), anti-GSDMD (209845, Abcam), anti-GAPDH (G8795, Sigma).

Cell lines culture and transfection

BV2 cells and HEK293T cells (ATCC) were maintained in Dulbecco's modified eagle media (DMEM; Corning) supplemented with 10% heat-inactivated fetal bovine serum (Biowest), 1% pen/strep (Corning), 2 mM L-glutamine (Corning), and 1% HEPES (Corning). To obtain primary bone-marrow-derived macrophages (BMDMs), bone marrow cells were collected from mouse femurs and tibia. Cells were then cultured and differentiated in non-tissue culture treated dishes with BMDM media (DMEM, 10% heat-inactivated fetal bovine serum, 10% CMG14 conditioned media, 2 mM L-glutamine, 1% pen/strep and 1% HEPES) and incubated for seven days at 37°C and 5% $CO_2$. All cell lines were mycoplasma negative. Transient transfection of HEK293T cells was performed using the Lipofectamine 3000 (Invitrogen) according to the manufacturer's instructions.

Mouse strains and infections

C57BL/6J, *Casp1/11*$^{-/-}$ (Jackson Laboratories) and *Stat1*$^{-/-}$ (Jackson Laboratories)(*40*) mice in this study were bred and maintained under pathogen-free conditions in the animal care facility at UT Southwestern Medical Center. *Ripk3*$^{-/-}$ mice were kindly provided by Z.Wang (UT Southwestern Medical Center). Mice were used for infections between 6-10 weeks of age. Mice were perorally (PO) inoculated with 25 µl of $10^6$ PFU MNoV diluted in D10 (DMEM with 10% FBS). Single fecal pellet and tissues were harvested into 2 ml tubes with 1mm zirconia/silica beads (Biospec). All of samples frozen in a dry ice/EtOH bath and stored at -$80^0$C. All experiments were performed according to experimental protocols approved by the Institutional Animal Care and Use Committee and complied with all relevant ethical regulations.

Generation of viral stocks and in vitro MNoV infections

MNoV$^{CW3}$ (accession EF014462.1) and MNoV$^{CR6}$ (accession JQ237823) were generated from plasmids (*41*). Briefly, plasmids encoding viral genomes were transfected into HEK293T cells to generate P0 stock. Cells were frozen at 48 hours post-transfection. The P0 virus were expanded by two passages in BV2 cells at MOI=0.05 to generate a P2 stock. To generate viral stocks, infected BV2 cells were freeze/thawed, cell lysate and supernatant were pelleted at 1200 g for 5 minutes, supernatant was filtered through a 0.22 µM filter and concentrated with 100,000 MWCO Amicon Ultra filter. Virus stocks were aliquoted, frozen at -80C and titered via plaque assay. MNoV$^{CW3}$ΔN, MNoV$^{CR6}$ΔN, MNoV$^{CW3}$ΔN20, MNoV$^{CR6}$ΔN20 were generated by PCR from parental pCR6 and pCW3 plasmids. The PCR products were gel purified and assembled using Gibson Assembly Master Mix (NEB). All plasmids were verified by DNA sequencing. Mutant viruses were propagated as for wildtype MNoV$^{CW3}$and MNoV$^{CR6}$, except P0 virus were expanded by two passages in BV2 and NTPase inducible cell line (1:1). Doxycycline was added to media after 12 h infection to induce cells death and release the virus. This was done for multiple rounds of infection to achieve high yield of virus particle. The primers for mutant viruses: MNoV$^{CW3}$ΔN20 Forward primer: GAATGGCAGGCCGAAGGGCCCTTTGGCCTCACCTCTGA; MNoV$^{CW3}$ΔN20 Reverse primer: TCATAGACCTTGTTCTGGAGGCCGAAATCATCA; MNoV$^{CW3}$ΔN Forward primer: GAATGGCAGGCCGAAGGGCTTGATGAGGAGGAGCA; MNoV$^{CW3}$ΔN Reverse primer: TCATAGACCTTGTTCTGGAGGCCGAAATCATCA; MNoV$^{CW3}$ Plasmid Forward primer: AACAAGGTCTATGACTTTGATGCCG MNoV$^{CW3}$ Plasmid Reverse primer: CCCTTCGGCCTGCCAT; MNoV$^{CR6}$ΔN20 Forward primer: AGTGGCAGGCTGAGGGTTTTGGCCTGACATC; MNoV$^{CR6}$ΔN20 Reverse primer: GTCATAGGTCTTGCTTTGGAGGCCGAAGTC; MNoV$^{CR6}$ΔN Forward primer: AGTGGCAGGCTGAGGGTCTTGATGAAGAAGAG; MNoV$^{CR6}$ΔN Reverse primer: GTCATAGGTCTTGCTTTGGAGGCCGAAGTC; MNoV$^{CW6}$ Plasmid Forward primer: AGCAAGACCTATGACTTTGAT; MNoV$^{CW6}$ Plasmid Reverse primer: ACCCTCAGCCTGCCACTCTCCAA.

BV2 and BMDMs infected with MNoV strains at indicated MOI for 1 hour. Viral inoculum was removed and washed with PBS twice and then media was added to the cells (*42*).

Virus quantification by plaque assay and Quantitative PCR

BV2 cells were seeded at 2 × 10$^6$ cells/well in 6-well plates. Plaque assay was performed by freeze/thawing infected samples followed by ten-fold serial dilutions. Media was removed from the BV2 cells and infected with serial dilutions samples and gently rocked for 1 hour. After inoculum was removed and cells were overlaid with 2 mL of overlay media (MEM containing 1% methylcellulose and 10% FBS). Plates were incubated for 48 hours prior to plaque visualization with crystal violet solution (20% ethanol/ 0.2% crystal violet).

MNoV genome copies in fecal pellets and tissues were determined as previously described (*43*, *44*). Briefly, RNA was isolated from fecal pellets using the ZR Viral RNA Kits according to manufacturer's protocol (Zymo Research). Tissue RNA extraction was performed using TRIzol (Sigma-Aldrich) and purified using RNeasy Mini Kit (Qiagen) according to manufacturer's instructions. cDNA was generated with SuperScript VILO cDNA Synthesis Kit (Thermo Fisher Scientific). Then, TaqMan assays was performed for the samples and standard curves using MNoV specific oligonucleotides: Probe: 5' FAM-CGCTTTGGAACAATG-MGBNFQ 3'; Forward primer: 5' CACGCCACCGATCTGTTCTG 3'; Reverse primer: 5' GCGCTGCGCCATCACTC 3'. A standard generated using RNA from ten-fold series dilution of parental virus stocks was used to quantitate viral genome equivalents in genome copy from mutant virus stocks. For measure the titer of mutant virus stocks, viral RNA was extracted from the 10-fold serial dilution of mutant virus stocks and parental virus stocks which known plaque-forming units (PFU). Ten-fold serial dilutions ranging from $10^6$ to 10 copies of standard plasmid was used for quantification of viral genome copy numbers. The plaque-forming units (PFU) of mutant virus stocks was calculated based on the equivalent genome copy of parental virus stocks.

Lentivirus production and transduction

For lentivirus production, lentivirus harboring the desired genes together with the packing plasmids pSPAX2 and pMD2.G were transfected into HEK293T cells by Lipofectamine 3000 (Invitrogen). The transfection medium was replaced with fresh culture medium (DMEM with 10% FBS) after 6 h transfection. Supernatants were collected at 48 h, cleared by centrifugation at 1,500 rpm for 5 min and filtered through 0.45 µM membrane (Millipore). For lentivirus transduction, 4 × $10^5$ BV2 cells were seeded into 6-well plates. Cells were transduced with the indicated lentivirus in transduction medium (DMEM with 10% FBS and 8 µg/ml polybrene) by spinning at 1,000g for 30 min at 37 °C. The transduction medium was replaced with culture medium after 6 h and selected with 2.5 µg/ml of puromycin (Sigma-Aldrich) and 50 µg/ml hygromycin (Invitrogen) after 48 h transduction. Viruses for the pLVX-Tet3G and pLVX-TRE3G-NTPase were generated by transfecting HEK293T cells using the Lipofectamine 3000 (Invitrogen). BV2 cells were co-infected with the pLVX-Tet3G and pLVX-TRE3G-NTPase viruses and selected with G418 (400 µg/ml) and puromycin (2.5 µg/ml). For the induction of NTPase expression, doxycycline was added to the tetracycline free media at a concentration of 2 µg/ml.

Generation knockout cell lines by CRISPR/Cas9

Guide RNA was cloned into lenti-CRISPR v2 puro and lenti-CRISPR v2 hygro vector (Addgene 98290 and 98291) according to the manufacturer's protocol (*45*). Lentivirus was produced as described above. 4 × $10^5$ BV2 cells were transduced with the indicated lentivirus and incubated for 48 h. Transduced cells were then selected with 2.5 µg/ml of puromycin (Sigma-

Aldrich) and 50 μg/ml hygromycin (Invitrogen). Single colony cells were then isolated and screened by western blot analysis for protein expression or determined by DNA sequencing. sgRNA sequences for targeting the genes of interest were as follows: *Caspase3* (AATGTCATCTCGCTCTGGTA), *Caspase9* (CACACGCACGGGCTCCAACT), *Ninj1* (TGCCAACAAGAAGAGCGCTG), *Bak* (ATCTTGGTGAAGAGTTCGT), *Bax* (CTGAACAGATCATGAAGAC), *Mlkl* (CTCATTCACTTCATGGAAG), *Gsdmd* (AGCATCCTGGCATTCCGAG).

Cytotoxicity assays

Cell death was quantitated by assaying the activity of LDH released into cell culture supernatants after infection and transfections using the CytoTox 96 Non-Radioactive Cytotoxicity Assay kit (Promega) according to the manufacturer's protocol. Cell viability was determined by measuring ATP levels using the CellTiter-Glo Luminescent Cell Viability Assay (Promega) according to the manufacturer's instructions.

IncuCyte live-cell imaging

For IncuCyte analysis, cells were seeded in 24-well plates. After overnight incubation, cells infected with MNoV or treated with doxycycline, and then 25 nM SytoxGreen (Invitrogen) were added. The plates were moved into an IncuCyte live cell imaging system. Cells were imaged every 30 minute and the SytoxGreen labeled cells (counted as dead cells) were quantified by the IncuCyte Zoom software.

Flow cytometric analyses

To measure cell death after MNoV infection, cells were stained with FITC Annexin V Apoptosis Detection Kit (BioLegend) according to the manufacturer's instructions. To assess mitochondrial membrane potential, cells were stained with the MitoTracker Green and MitoTracker Deep Red (50 nM each) or TMRM (50 nM) for 30 min at 37°C. To assess mitochondrial ROS production, cells were stained with the mitochondria-specific superoxide indicator MitoSox Red for 30 min at 37°C. Cells were then washed with phosphate buffered saline (PBS) and analyzed using BD FACS Calibur.

Mitochondria isolation and cytochrome *c* release assay

Mitochondria were purified from stable cell lines using the mitochondrial isolation kit (Abcam, ab110168) according to the manufacturer's instructions. The cytosolic fraction and mitochondrial fraction were collected for western blot. For cytochrome *c* release assay, BV2 cells were collected and washed in phosphate-buffered saline (PBS) and resuspended in 5 times volume of isotonic buffer (10 mM Tris-Cl pH 7.5, 10 mM KCl, 250 mM sucrose, 1.5 mM $MgCl_2$) and incubated on ice for 15 min. Cells were then passage through a 22-gauge needle 25 times and were centrifuged at 1,000 g for 10 min at 4°C to pellet nuclei. The supernatant was collected and centrifuged at 7,000 g for 10 min at 4°C to collect mitochondria. Isolated mitochondria were incubated with recombinant NTPase (5 μM) for 30 min at 37°C. After treatment, mitochondria were pelleted by centrifugation (7,000 g, 4°C, 10 min) and supernatant was collected. The pelleted mitochondria were lysed in lysis buffer (50 mM Tris-Cl (pH 7.4), 150 mM NaCl) supplemented with 1 mM EDTA, 1% Triton X-100, 0.1% SDS, 0.5% deoxycholate and protease inhibitor cocktail (Roche), followed by centrifugation (15,000 g, 4°C, 10 min). Then, SDS loading buffer was added to each fraction and analyzed with western blot.

Purification of recombinant proteins

To obtain full-length, N-terminal and C-terminal NTPase proteins, *E. coli* BL21cells harboring the indicated plasmid (pMAL-c6T × $His_6$-MBP vector) were grown in LB medium. Protein expression was induced overnight at 20°C with 0.3 mM isopropyl β-D-1-thiogalactopyranoside (IPTG) after OD600 reached 0.8. Cells were harvested by centrifugation and lysed in the buffer containing 20 mM Tris-HCl (pH 8.0), 300 mM NaCl and 10 mM β-mercaptoethanol followed by sonication. The recombinant protein was affinity-purified by amylose resin (NEB) and eluted with buffer containing 20 mM Tris-HCl (pH 8.0), 300 mM NaCl, 10 mM β-mercaptoethanol and 10 mM maltose and further purified by Superdex 200 gel-filtration column (GE Healthcare). For liposome leakage assay, the $His_6$-MBP tag was removed by overnight TEV digestion at 16°C and further purified and concentrated by Superdex 200 gel-filtration column (GE Healthcare).

Immunoblot and immunoprecipitation

Cells were lysed in lysis buffer (50 mM Tris-Cl (pH 7.4), 150 mM NaCl) supplemented with 1 mM EDTA, 1% Triton X-100, 0.1% SDS, 0.5% deoxycholate and protease inhibitor cocktail (Roche) and then spun down at 15,000 g for 10 mins. The supernatants were boiled in SDS loading buffer for 10 min before electrophoresis with 4-12% Bis-Tris plus gels (Thermo Fisher Scientific). The resolved proteins were then transferred to polyvinylidene difluoride (PVDF) membrane (Millipore), which was probed with the indicated antibodies. Protein bands were developed using Luminata Forte Western HRP substrate (Millipore) or Super Signal West Pico chemiluminescence ECL kit (Pierce). For immunoprecipitations, cell extracts were prepared using lysis buffer (50 mM Tris-HCl (pH 7.4), 150 mM NaCl, 1 mM EDTA, 1% Triton X-100, 0.1% SDS, 0.5% deoxycholate) containing complete protease inhibitor cocktail (Roche). Lysates were incubated with FLAG-M2 agarose beads (Sigma-Aldrich) for 4 hours at 4 °C. After five times washes with lysis buffer, proteins bound to FLAG-M2 agarose beads were eluted using 3 × FLAG peptides (Sigma-Aldrich), followed by immunoblot analysis.

Immunostaining and confocal microscopy

Cells were grown on coverslips, fixed with 4% paraformaldehyde in PBS for 15 min, and permeabilized for 30 min in 0.1% Triton X-100 in PBS and blocked using 5% BSA for one hour at room temperature. Then, the cells were stained with the indicated primary antibodies, followed by incubation with Alexa Fluor secondary antibodies (Life Technologies). Nuclei were stained with DAPI (Cell Signaling Technology). Images were captured using a confocal microscope (Zeiss LSM 700 confocal microscope) with a 63 × oil immersion objective.

Protein-lipid binding assay

The protein-lipid binding assay performed with on Membrane Lipid Strips (Echelon Biosciences) according to the manufacturer's instructions. Lipid strips blocked with binding assay buffer (3% fatty acid-free BSA in PBS) for 1 h and incubated with protein (1 μg/ml) diluted in binding assay buffer for 1 h and then washed three times with wash buffer (0.1% Tween-20 in PBS). Membrane-bound proteins were then detected by immunoblot.

Liposome preparation

Natural and synthetic lipids were dissolved in chloroform. Lipids with indicated compositions were mixed in a glass vial. The lipid films were obtained by evaporating under a stream of nitrogen and the dry lipid film was then hydrated at room temperature with constant mixing in 500 μl buffer A (20 mM HEPES (pH 7.5) and 150 mM NaCl). Liposomes were generated by extrusion of the hydrated lipids through 100 nm polycarbonate filter (Whatman) 20 times using the Mini-Extruder device (Avanti Polar Lipids Inc.). To prepare $Tb^{3+}$-encapsulated liposomes, the lipid film was hydrated with 500 μl buffer B (20 mM HEPES (pH 7.5), 100 mM NaCl, 50 mM sodium citrate and 15 mM $TbCl_3$). After the extrusion process, $Tb^{3+}$ ions outside the liposome were removed by washing with buffer A on a centrifugal filter device (Amico Ultra-4, 100K MWCO, Millipore). The liposomes were subjected to buffer A for use. All the liposomes were stored at 4°C and used within 48 h.

Liposome binding and leakage assay

For liposome binding assay, the indicated proteins (0.2 μM) were incubated with liposomes at room temperature for 30 min in 0.1mL of buffer A. The liposomes were pelleted by centrifugation in a Beckman Optima XE-90 Ultracentrifuge at 40,000 rpm for 20 min at 4 °C. The supernatants (S) were collected, and the pellets (P) were washed twice with buffer A and then resuspended in same volume as supernatants. Proteins in both supernatant and pellets were then analyzed by SDS-PAGE and immunoblot. For liposome leakage assay, $Tb^{3+}$-entrapped liposomes were suspended in 100 μl buffer A supplemented with 50 μM of DPA and 0.5 μM indicated NTPase recombinant proteins were added. The emission fluorescence at 490 nm after excitation at 276 nm as $F_t$ was continuously recorded for 10 min at 20 s intervals using a Tecan Infinite M200 Pro plate reader. At the end of the incubation, 10 μl of 1% Triton X-100 was added to measure complete release of $Tb^{3+}$. The percentage of liposome leakage is defined as: leakage ($t$) (%) = $100 \times ((F_t - F_{t0})/ (F_{t100} - F_{t0}))$, where $F_{t0}$ is the emission fluorescence of the $Tb^{3+}$ liposomes before proteins were added, $F_t$ is the fluorescence reads recorded at individual time points, and $F_{t100}$ is mean values of the top three fluorescence signal after adding 0.1% Triton X-100

Protein modeling and phylogenetic analysis

Sequences were retrieved from the NCBI database. Accession numbers and information related to the sequences are present in supplemental table 1. Representative sequences for caliciviruses were selected primarily based on exemplar isolates from ICTV for the individual genera. (https://talk.ictvonline.org/ictv-reports/ictv_online_report/positive-sense-rna-viruses/w/caliciviridae). Unclassified species from ICTV for individual calicivirus genera were included to increase phylogenetic sampling. Mouse MLKL domains are annotated based on the predicted structure from (PDB: 4BTF) (*38*, *46*). Amino acids sequences were analyzed, aligned using MUSCLE, and visualized with Geneious Prime. Presentation of sequences is based on known host and viral species relationships (*47*). Expanded alignments with additional host and viral species are present in supplemental table 1.
